# ASM Variants in the Spotlight: A Structure-Based Atlas for Unraveling Pathogenic Mechanisms in Lysosomal Acid Sphingomyelinase

**DOI:** 10.1101/2023.11.24.568551

**Authors:** Simone Scrima, Matteo Lambrughi, Matteo Tiberti, Elisa Fadda, Elena Papaleo

**Affiliations:** Cancer Structural Biology, Danish Cancer Institute, 2100, Copenhagen, Denmark; Cancer Systems Biology, Section for Bioinformatics, Department of Health and Technology, Technical University of Denmark, 2800, Lyngby, Denmark; Department of Chemistry and Hamilton Institute, Maynooth University, Maynooth, co. Kildare, Ireland

**Keywords:** variants of uncertain significance, acid sphingomyelinase deficiency, molecular dynamics simulations, protein structure, variant interpretation

## Abstract

Lysosomal acid sphingomyelinase (ASM), a critical enzyme in lipid metabolism encoded by the SMPD1 gene, plays a crucial role in sphingomyelin hydrolysis in lysosomes. ASM deficiency leads to acid sphingomyelinase deficiency, a rare genetic disorder with diverse clinical manifestations, and the protein can be found mutated in other diseases. We employed a structure-based framework to comprehensively understand the functional implications of ASM variants, integrating pathogenicity predictions with molecular insights derived from molecular dynamics simulations in a lysosomal membrane environment. Our analysis, encompassing over 400 variants, establishes a structural atlas of missense variants of lysosomal ASM, associating mechanistic indicators with pathogenic potential. Our study highlights variants that influence structural stability or exert local and long-range effects at functional sites. To validate our predictions, we compared them to available experimental data on residual catalytic activity in 135 ASM variants. Notably, our findings also suggest applications of the resulting data for identifying cases suited for enzyme replacement therapy. This comprehensive approach enhances the understanding of ASM variants and provides valuable insights for potential therapeutic interventions.

## Introduction

Acid sphingomyelinase (ASM), encoded by the SMPD1 gene, is a mammalian phosphodiesterase found in lysosomes and in the extracellular space. It plays a critical role in lipid metabolism by hydrolyzing sphingomyelin to ceramide and phosphocholine^1^. The ASM precursor is a 75 kDa pre-protein that undergoes post-translational processing, generating both the extracellular secreted and the 70KDa lysosomal (l-ASM, hereby called ASM) forms^1,2^. Lysosomal ASM, with its structural complexity and tight regulation, plays an important role in cellular processes, including immune response, endocytosis, and plasma membrane repair^3,4^.

Structurally, ASM comprises four domains: i) the saposin domain (residues 86-169), ii) the proline-rich linker (residues 170-197), iii) the catalytic metallophosphatase domain (residues 198-540), and iv) a helical C-terminal domain (CTD, residues 541-613)^5,6^ (**Figure 1A**). The experimental structures of ASM reveal functionally related conformational changes in the saposin domain, which can adopt both open and closed states around the active site^5,6^. In the closed state, the saposin domain interacts with the CTD, forming a compact globular structure around the active site. In the open state, the saposin domain exhibits a V-shaped conformation, exposing a concave hydrophobic surface toward the active site to accommodate the hydrophobic ceramide chains of sphingomyelin. Additionally, the proline-rich linker is a rigid region, wrapping around the catalytic domain and assuming an L-shaped strap conformation ^5,6^. Furthermore, the catalytically active form of ASM is predominantly monomeric^7^, despite suggested dimeric states.^8^

**Figure 1.**
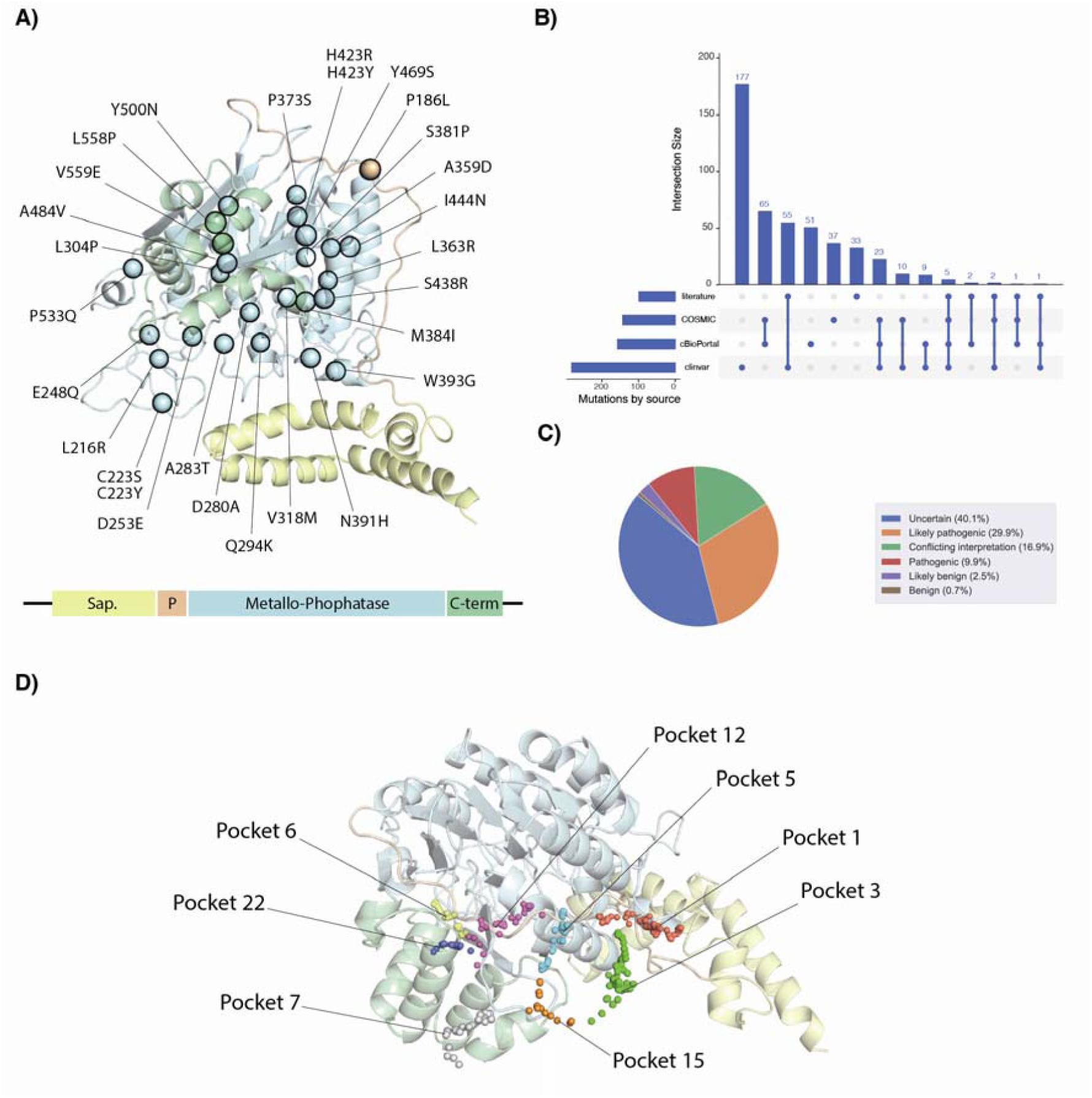
ASM structure, variants, and functional interfaces. A) Cartoon representation of ASM_86-613_ (PDB ID: 5I85) with spheres highlighting the C= atom of the 26 residues where 28 pathogenic variants have been found in the ClinVar database. Below is the structure of the diagram of ASM domains: in yellow saposin, orange proline-rich linker, cyan the catalytic metallophosphatase, and green C-terminal domain. The Zn^2+^, glycosylations, and phosphocholine in the X-ray structure were omitted for clarity of view. B) Upset plot with an overview of the collected variants and their overlap across the different sources used in the study. C) Overview of all the variants collected from the ClinVar database, their classification, and percentages, using the internal MAVISp interpretation of the ClinVar data. D) Pockets on the surface of the protein structure as identified by Fpocket. We visualized only the pockets with residues connected either with the protein active site or associated with the membrane according to the analysis of the MD simulation.

The catalytic site includes two Zn^2+^ ions with a trigonal bipyramidal coordination geometry. The Zn^2+^ ions are coordinated by four histidines (H210, H427, H459, H461), two aspartates (D208, D280), one asparagine (N320), and a catalytic water molecule^5^ (**Figure 1A**). Moreover, N327, E390, and Y490 have been suggested to be involved in substrate binding^5^. According to the proposed catalytic mechanism of ASM, the Zn^2+^ ions may also play a role in substrate recognition by anchoring the phosphate oxygens of the headgroup of sphingomyelin. Furthermore, it has been suggested that the active site geometry, especially H321 or H284, facilitates proton donation to the oxyanion of the ceramide leaving group^5^. The phospholipid cleaving activity of ASM is also regulated by membrane lipids such as anionic bis(monoacylglycero)phosphate found in late endosomes/lysosomes and interactions with other lipid-binding proteins^9^. The mature form of ASM includes six N-glycosylation sites at the residues N86, N175, N335, N395, N503, and N520 ^5,10^. Moreover, the structure features eight disulfide bridges, including C91-C167, C94-C159, C122-C133, C223-C228, C229-C252, C387-C433, C586-C590, and C596-C609 ^11^.

ASM deficiency leads to acid sphingomyelinase deficiency (ASMD), a rare genetic disorder^12,13^. ASMD results in the accumulation of sphingomyelin and other molecules in various tissues, with varying levels of severity. The spectrum ranges from a fatal neurodegenerative disorder in children (Niemann-Pick disease type A, NPDA) to milder forms in adults (Niemann-Pick disease type B, NPDB)^14^. ASMD is primarily caused by mutations in the SMPD1 gene, inherited in an autosomal recessive manner. Common challenges in managing ASMD include misdiagnosis and delayed care due to the paucity of consensus clinical guidelines^12^.

As with many other disease-related genes, the volume of data on variants found in patients is growing, accompanied by an increasing number reported as variants of uncertain significance (VUS) or with conflicting evidence regarding their functional effects. This underscores the need for a comprehensive understanding of ASM variants and their functional implications. Recently, an approach based on different sequence and structural features has been used to classify ASM variants^15^. In another study, one of the recent models for predicting pathogenicity based on evolutionary-related properties, i.e., EVE^16^ was applied to study a group of variants from SMPD1^17^.

However, the aforementioned studies do not directly link to the underlying molecular mechanisms exerted by the mutations. Understanding these mechanisms is crucial for devising strategies to overcome variant effects and design effective treatments. In this context, we introduced a structure-based framework, i.e., MAVISp (Multi-layered Assessment of VarIants by Structure for proteins)^18^ to integrate pathogenicity prediction with a molecular understanding of the effects of disease-related mutations. To address the gap of knowledge regarding functional changes induced by ASM variants, we thus applied MAVISp. This was done in conjunction with the pathogenicity scores included in the framework^16,19,20^, as well as the analysis of different properties from all-atom explicit solvent molecular dynamics (MD) simulations of ASM in a membrane environment mimicking the lysosomal compartment to study the effect of missense variants in ASM. Our study also provided the introduction of new modules for variant interpretation within MAVISp, covering local and long-range effects on metal binding regions and active sites. Focusing on more than 400 variants identified by ClinVar^21,22^, literature-based curation or in cancer samples, we aimed to provide the first comprehensive atlas of the effects of missense variants on lysosomal ASM at the structural level. We identified 31 variants for which we could associate a mechanistic indicator to their pathogenic potential, integrating MAVISp data, analysis of MD simulations, pathogenicity scores, and experimental data on ASM activity. Furthermore, we carried out in silico saturation mutagenesis, assessing a total of 10560 variants and identified more than 1500 potentially pathogenic variants with at least a linked mechanistic indicator.

## Results and Discussion

### Overview of the variants included in the study

We analyzed 471 missense variants in ASM (residues 86-613), sourced from ClinVar^21,22^ (282), cBioPortal^23^ (157), and COSMIC^24,25^ (143) as illustrated in **Figure 1B**. Additionally, we included 109 variants from the literature (**Figure 1B** and **Table S1**). The literature-based curation partially overlapped with the data retrieved from the databases mentioned above. Specifically, 55 variants are also present in ClinVar, nine in both COSMIC and cBioPortal and five across all the databases (**Figure 1B**). Of the collected variants, 67% reside in the ASM catalytic domain, while the remaining are located in the saposin domain (14%), the proline-rich linker (5%), and the C-terminal domain (14%). This distribution aligns with the current knowledge that most Niemann-Pick A and B variants occur within the ASM catalytic domain^5^. Approximately 40% of the variants are classified as Variants of Uncertain Significance (VUS) as can be seen in **Figure 1C**. Among the ClinVar variants, eight are reported with pathogenic effects and two are classified as benign.

### Overview of the structural analysis included in the study using the MAVISp framework

In our study, we focused on the open state of ASM, which represents the conformational state important for the protein activity and membrane association^5,6^. Moreover, no experimental structures are available for the closed state of the human variant. While X-ray structures of the ASM homodimer were reported, ASM predominantly exists as a monomer in solution^5,7^. Consequently, our study specifically concentrated on the monomeric ASM structure, using an X-ray structure solved at a resolution of 2.5 Å^5^. We used the structure to perform the calculations covered by the simple mode of MAVISp^18^, which relies on the usage of one single conformation for the protein of interest. Additionally, we performed a one-μs MD simulation of the N-glycosylated variant of the protein associated with the membrane for the ensemble mode of the MAVISp framework. In the MD simulation, we considered a protonation state for the protein residues within the pH range observed in lysosomes, i.e., 4.5-5.0^5^.

MAVISp, with its INTERACTOME module^18^, facilitates the identification of protein-protein complexes for the structural studies of the effects of mutations on interactions. We identified six potential interactors for ASM with Mentha scores below 0.5 (**Table 1**). CASP7, with the highest score, was prioritized considering its known role in ASM cleavage at the D253-L254 site during bacterial infection and membrane damage, contributing to membrane repair^26–28^. Unfortunately, attempts to model this interaction using AlphaFold Multimer^29^ were unsuccessful (**Figure S1**), leading to the exclusion of CASP7 from further binding free energy calculations. In addition, the remaining interactors in **Table 1** were excluded due to insufficient literature-based support for a direct association with SMPD1.

**Table 1.** Protein interactors for ASM. The results from the Mentha2PDB workflow of MAVISp are reported in the table with their associated Uniprot ID, gene name, Mentha score, and PMID. ‘AFmulti’ indicates a case in which we generate a model with AF-Multimer. ‘QC’ indicates the Quality Control step within the MAVISp framework for selecting protein-protein complexes.

| Interactor Uniprot ID | Interactor Gene name | Metha score | PDB structure | PMID |
| --- | --- | --- | --- | --- |
| P55210 | CASP7 | 0.454 | AFmulti - not passed QC | 21157428 |
| P20073 | ANXA7 | 0.332 | N.A. | 21900206 |
| P09525 | ANXA4 | 0.332 | N.A. | 21900206 |
| P55055 | NR1H2 | 0.332 | N.A. | 21900206 |
| Q9BVJ7 | DUSP23 | 0.332 | N.A. | 21900206 |
| Q14790 | CASP8 | 0.309 | N.A. | 21157428 |

Considering the results above, we decided to retain for the study specific structural descriptors from the MAVISp framework. These include: i) effects on folding free energy (i.e., stability of the protein architecture), ii) effects due to interplay with phosphorylations, iii) effects transmitted long-range to solvent-exposed interfaces, to the Zn^2+^-binding site or the residues essential for catalysis (H321 or H284) and substrate recognition (N327, E390 and Y490), along with iv) local effects on the active site and the Zn^2+^-binding region.

In connection with the prediction of long-range effects, the MAVISp framework applies a method to select potentially relevant interfaces as response sites from the raw allosteric map of AlloSigMA 2 ^30^ based on pocket identification. We identified 24 distinct pockets in the X-ray structure of ASM (see OSF repository). After refining the selection, eight pockets were identified involving residues for substrate binding, catalytic function, or membrane association (**Figure 1D**). Specifically, pockets 1, 3, 6, 7, 15, and 22 refer to regions important for interactions with the membrane (according to the results of the MD simulation reported in this study). In contrast, pockets 5 and 12 were in the proximity to the catalytic site.

Additionally, we implemented new modules in the MAVISp framework to assess the influence of the mutations on binding to the metals or on the catalytic activity in terms of local and long-range effects (see Materials and Methods).

### Analysis of variants reported as pathogenic or benign in ClinVar

At first, we focused on the variants reported with a clear pathogenic or benign interpretation in ClinVar to evaluate if the modules used in MAVISp can shed light on the mechanisms underlying their effects as well as to evaluate the accuracy of the pathogenicity scores used by the framework. To this goal, we analyzed the 27 pathogenic and 2 benign variants reported in ClinVar **(Table S2, Figure 2A**). We noticed that at least two out of three pathogenicity scores used in the MAVISp framework identify as damaging a total of 21 variants reported as pathogenic in ClinVar (C223S, C223Y, D253A, D280A, A283T, Q294K, A359D, L363R, S381P, M384I, N391H, W393G, H423R, H423Y, S438R, I444N, Y469S, A484V, Y500N, L558P, and V559E). AlphaMissense^19^ and DeMaSk^20^ agree on a neutral effect for the benign variants reported in Clinvar, i.e., P187S and G508R. On the contrary, EVE^16^ results in an uncertain prediction for P187S, G508R, and other variants, primarily due to inadequate coverage within the multiple sequence alignment used in this study (see OSF repository). Overall, AlphaMissense^19^ reported as damaging variants all the variants reported in ClinVar with a pathogenic effect with some exceptions. All the cases with AlphaMissense reported a ClinVar pathogenic variant as benign corresponded to cases where the other pathogenicity scores also failed to identify a damaging effect. In detail, V318M and P533Q were reported as uncertain from AlphaMissense. The structural analysis carried out in MAVISp did not identify a mechanistic indicator (i.e., a structural effect) associated with these variants. Moreover, V318M is reported with a ClinVar review status of 0, and caution should be taken to report it as a genuinely pathogenic variant without solid evidence. In the study reported in ClinVar, V318M was identified in a patient with a severe Niemann-Pick disease type A phenotype, showing heterozygous missense variants V318M and Y500N^31^. Y500N is reported as pathogenic in ClinVar and predicted damaging by AlphaMissense. MAVISp associates it with a destabilizing effect due to changes in structural stability (**Figure 2A**). The severe phenotype observed might be attributed primarily to Y500N. At the same time, V318M might have a marginal effect and could be reclassified as benign after further experimental validation. Additionally, we could rule out an effect on ASM activity. Indeed, V318 is located within the β3-α3 loop and its side chain is oriented towards the aromatic and hydrophobic residues of α3 and α4, unlikely to alter the active site (**Figure S2**). P533Q is reported as associated with Niemann-Pick disease type A and a ClinVar review status of 1. We also identify this variant as a somatic one in cancer samples, where most mutations are expected to be passenger and neutral^32^. Fluorescence-based enzymatic activity assay reported that another variant at P533 (P533L, reported in ClinVar with conflicting interpretation) causes a very mild reduction of ASM activity (i.e., > 70% residual activity, **Table S3**^33^) suggesting that this variant is unlikely to have a damaging effect on the protein function. Nevertheless, we cannot rule out mild effects on structural stability since the simple and ensemble modes of MAVISp provided an uncertain classification based on changes in folding free energy upon mutation.

**Figure 2.**
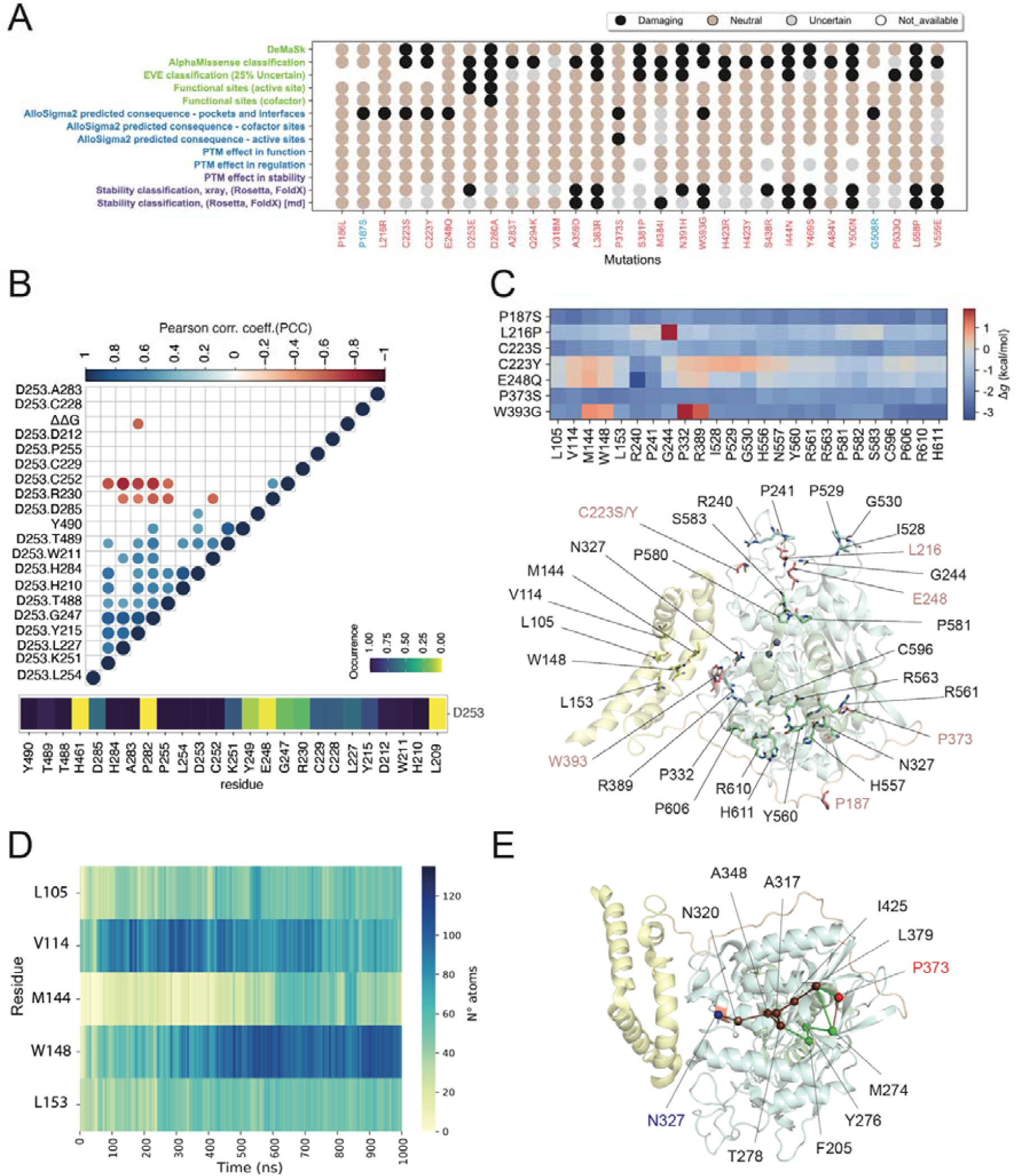
Analysis of pathogenic and benign variants of ASM reported in ClinVar. A) The dot plot from the downstream analysis toolkit of MAVISp illustrates the link between the predicted pathogenic effect and the different mechanisms at play, as covered by the MAVISp framework. We highlighted the known benign and pathogenic variants from ClinVar in light blue and black, respectively. B) Correlation analysis. The upper figure represents the Pearson correlation for each pair of properties analyzed for the mutation site D253. Additional measurements can be found in Figure S2. Notably, the changes in folding free energy are tightly associated with changes in the contacts between D253 and Y215. The lower panel includes a heatmap with the occurrence of contacts for D253 and its surrounding residues in the MD simulation. C) The heatmap shows the results of predicted long-range effects for the variants P187S, L216P, C223S/Y, E248Q, P373S, and W393G, as predicted by the LONG RANGE module of MAVISp. The mutation sites and their response sites are also visualized on the 3D structure of ASM in the lower panel. D) The heatmap illustrates the contacts with lipids in the MD simulation for the distal response sites L105, V114, M144, W148, and L153. E) The paths of communication to the response site N327 (in blue) for the mutation site P187 (in red) are reported with the nodes represented by spheres centered on the Cα atoms of the residues and the edges as cylinders. The paths have been calculated from an atomic contact PSN based on the interaction strength and occurrence in the MD simulation.

Next, we focused on the mechanistic indicators proposed by MAVISp to explain the damaging impact of the ClinVar pathogenic variants.

D280A pathogenic effect is associated with alteration of the catalytic activity and Zn^2+^-binding (**Figure 2A**). D280 belongs to the Zn^2+^-binding site and directly interacts with the proposed catalytic residue H321, in agreement with an abolishment of the catalytic activity as measured experimentally^34^.

MAVISp associates a destabilizing effect on structural stability to nine of the pathogenic ClinVar variants (A359D, L363R, M384I, W393G, I444N, Y469S, Y500N, L558P, and V559E) with the ensemble mode (**Figure 2A**). The predicted changes in folding free energy for the A359D variant agree with previous results using different free energy-based algorithms on a model of the ASM protein structure^35^, where also a dramatic effect on the catalytic activity of the enzyme was measured (i.e., residual activity of 3.8%). L363R, M384I, Y469S were also reported with low residual activities (**Table S3**). W339G, L558P, Y500N, V559E, and I444N are suitable candidates to explore if these variants cause changes in protein degradation and activity in cellular assays or in assays to estimate ASM residual activity. Furthermore, three variants were reclassified from destabilizing in the simple mode to uncertain using the ensemble mode (i.e., D253E, N391H, and S438R). In the MAVISp ensemble mode, the reclassification of D253E from destabilizing to uncertain was due to the average ΔΔG values from FoldX and Rosetta falling in the uncertainty region, i.e., in the range of 2-3 kcal/mol threshold. On the other hand, for N391H and S438R, the change in classification is due to a discrepancy between FoldX and Rosetta predictions in the ensemble mode, with only FoldX (N391H) or Rosetta (S438R) predicting a destabilizing effect for the variant. These variants require further inspection for more conclusive data on their effects on structural stability. We thus investigated the interaction and conformational states of the three mutation sites in the MD simulation.

D253 and N391 are in the β1-α1 and β5-α5 loops of the catalytic domain of ASM, respectively, and define the wide-open lipid-binding cleft of ASM^5^ (**Figure 1A**). Part of the β1-α1 loop (i.e., the region 233-242) has high flexibility in the MD simulation, according to the analysis of the Cα Root Mean Square Fluctuation (**Figure S3**). In the structure of the simple mode, the side chain of D253 forms hydrogen bonds with the main chain and side chain of H284 (79% of MD frames) and the side chain of T488 (68%) (**Table S4**). We also observed that conformational changes in the loops where the residues are located affect these interactions and cause the breaking of the hydrogen bonds. To further explore the region, we employed a contact-based approach (see Methods) to assess the frequency of interactions between this residue and its neighbors. Indeed, the D253 side chain can form contacts with high occurrence with other residues located in the β1-α1, β2-α2, and β9-β10 loops (**Figure 2B**). Mutations at D253 could affect the enzyme’s catalytic activity since the residue interacts with one of the two proposed catalytic residues, i.e., H284 (**Figure 2A**). However, the specific variant found in ASM at the D253 site, e.g., D253E, might retain hydrogen bonds with H284. MD simulations of the mutated variant could help to validate the prediction. To further evaluate if any of the changes in the interactions mediated by D253 affected changes in the predicted folding ΔΔG, we performed a correlation analysis, including different structural features and the ΔΔG values in the 25 structures of the MD ensemble used for the FoldX calculation for the MAVISp framework (**Figure 2B**). We identified a negative correlation between the formation of side-chain contacts between D253 and Y215 and the folding ΔΔG values (**Figure 2B**). D253E is predicted with values close to the cutoffs in the simple mode (i.e., 4.43 and 3.09 kcal/mol, with FoldX or Rosetta, respectively). In contrast, in the MAVISp ensemble mode, the average values decrease to approximately 2 kcal/mol for both methods. This makes it challenging to ascertain the precise impact of this variant on structural stability. In particular, it has been shown that when ΔΔG values fall within the 2-3 kcal/mol range, the effects on cellular-level structural stability vary depending on the specific protein^36^, and experimental data would be needed to fine-tune a more specific threshold. From the analysis above, we can conclude that substituting aspartate with the longer side chain of glutamate causes steric hindrance only in conformations where Y215 is near the 253 site. According to the MD simulation, Y215 could be in different states thanks to the rearrangements of its sidechain. To better judge the impact of D253E, it would be important to estimate the populations of the different states, for example, using enhanced sampling approaches, as we did in p53^37^. Interestingly, D253E was reported in ClinVar from a single patient with neurodevelopmental delay and heterozygous with G168R. G168R is included in ClinVar as likely pathogenic, and AlphaMissense reports this variant as damaging. According to the MAVISp assessment, G168R can be associated with alterations in structural stability (see below). Thus, we could speculate that G168R could be the alteration with the main effects on the patient phenotype. N391H was reported in two infants showing early-onset symptoms classified as Niemann-Pick disease type A^38^. Furthermore, enzymatic activity assays revealed that N391H substantially decreases ASM activity, retaining only 2.61% of the activity observed in the wild type^39^. At the structural level, N391 is highly buried to the solvent (SASA LJ 0%), and its side chain forms hydrogen bonds with the main chain of F329 (46% in MD frames) and N385 (35% in MD frames, **Table S4**). Furthermore, N391 establishes stable contacts with various residues within the β3-α3 and β5-α5 loops (**Figure S4**). The results above and the experimental evidence suggest that the variant should be considered as altering structural stability.

Another variant reclassified from destabilizing to uncertain in the MAVISp ensemble mode, S438R, is located on α6 in the catalytic domain facing the C-terminal region of the protein (residues 600-611). In the structure of the simple mode, the side chain of S438 is fully buried and remains buried in the MD simulations (average SASA 3.22 %). In the MD simulation, we observed that S438 forms main chain hydrogen bonds with Y442 for almost half of the simulation (47% of MD frames) (**Table S4**). S438R introduces a positive charge and a bulkier residue but does not seem to lead to steric clashes in this region according to the analysis of the 125 models from FoldX for S438R shows that the arginine can assume different conformations, and in one of the scans is involved in a salt bridge with D604 (**see Supplementary Movie S1**). These results suggest that the effect of the variant is likely to be neutral for the structural stability of the protein. Takashi et al.^40^ proposed that, despite its pathogenic nature, the variant retains partial catalytic activity. This is supported by the fact that both patients in the study, who were homozygous for the mutation, exhibited symptoms consistent with Niemann-Pick type B and aligned with our result of no marked effects on structural properties.

Among the predicted pathogenic variants from AlphaMissense and at least one of the other pathogenicity scores, C223S/Y and W393G have been predicted with a long-range effect from the MAVISp framework. C223Y is reported with possible long-range effects on R240 according to the default assessment implemented in MAVISp (**Figure 2A**). From the structural point of view, a closer look suggests that it can be considered a false positive due to the proximity between the two residues. Conversely, the C223S and W393G variants featured stabilizing long-range effects on several distal response sites (**Figure 2A** and OSF repository). Due to the coarse-grain model applied in the long-range module and a workflow based on distance and energy cutoffs, it is good practice to explore further the results and validate them. To further validate the possible allosteric effects exerted by P373S and E248Q, we used a Protein Structure Network based on atomic contacts (PSN) applied to the MD simulations^41,42^, and we then estimated the shortest path of communication between C223S and W393G and the predicted response sites. We found no path linking the residues to their predicted response sites.

G508R and P187S are predicted as benign from pathogenicity scores and benign in ClinVar, but MAVISp identified a possible long-range effect. In the case of G508R, the assignment was due to the interaction with A195, which is in a loop facing G508 and is thus not considered a genuine long-range effect. P187S has predicted long-range effects on different distal sites according to Allosigma2, and we further verified through the PSN-MD approach if there were underlying communication paths. Still, no shortest paths were identified to any endpoint.

In addition, P373S, E248Q, P186L, and L216R were predicted as benign by AlphaMissense but reported pathogenic in ClinVar. For three of them (P373S, E248Q, and L216R), the long-range module of MAVISp identifies a potentially damaging effect. Indeed, all were predicted to have stabilizing long-range effects toward R240 and P241 on the β1-α1 loop that localizes near the H2-H3 turn within the saposin domain (**Figure 2C**). However, upon visual inspection of the 3D structure, we observed that the residues R240 and P241 are located near L216 and should not be considered a genuine signature of a long-range effect. Additionally, the P373S mutation is predicted to have a long-range stabilizing effect on different residues (i.e., L105, V114, M144, W148, L153, G244, N327, P332, R389, I528, P529, G530, and S583). W148, M144, L153, L105, and V114 are found within the saposin domain, delimiting pocket 1 (**Figure 1D**), and they all form contacts with the lipid bilayer during the MD simulation (**Figure 2D**). On the other hand, the remaining residues are located in the catalytic domain, with N327 and R389 located in pocket5, where the catalytic site is located. S583 is instead located in the C-terminal domain, in the pocket 22, which is one of the interfaces for interaction with the membrane. While we found no paths linking E248Q to the predicted endpoints from Allosigma2, we found three shortest paths linking P373 only to N327 of comparable path lengths, which could represent potential allosteric pathways (**Figure 2E, Table S5)**. This suggests that P373 is likely to be important as an end/starting point for communication to the catalytic domain where N327 is located.

E248Q is also reported with review status 1. Still, it is associated with Niemann-Pick A and B. Upon reviewing the available literature in ClinVar associated with this variant, the variant associated with the disease is reported to be E248K rather than the E248Q variant^43,44^. We based our numbering on the protein sequence from the NP_000534.4. RefSeq ID, which has 631 amino acids. This introduces a +2 amino acid shift compared to previously published positions (see Materials and Methods for more details). Thus, according to the reference sequence that we are applying E248K correspond to E246K. Nevertheless, an experimental study^45^ shows the importance of this variant by reporting that in vitro, it retains only partially catalytic activity (i.e., 30% residual activity, **Table S3**). On the other hand, L216R is associated with Niemann-Pick A. Although ClinVar does not report any specific studies on this variant, Ranganath et al. ^38^ suggests its relevance in the disease. This study identifies L216R as a novel variant, with the patient exhibiting a low ASM enzyme activity in leukocytes and a severe phenotype with developmental delay and failure to thrive. Moreover, Deshpande et al.^39^ demonstrates in vitro activity ranging from 0% to 2.6%. Our structural analyses did not identify mechanistic indicators for the effects of E248Q or L216R observed experimentally. It remains to be investigated if a residual activity of 30% for E248Q could cause a disease phenotype or exert a mild effect. On the other hand, P373S, despite being reported as pathogenic with review status 1 in ClinVar, has not been fully characterized with functional studies. Our results suggest a functional allosteric effect on the catalytic domain for future experimental studies.

### Analysis of variants reported as likely pathogenic or likely benign in ClinVar

Next, we analyzed the 85 likely pathogenic and seven likely benign variants reported in ClinVar (**Figure 3A-B**). We noticed that at least two of the three pathogenicity scores used in the MAVISp framework identify as damaging 65 variants reported as likely pathogenic in ClinVar. AlphaMissense classified the most likely pathogenic variants as damaging (73%) and the likely benign ones as neutral (**Figure 3A-B**).

**Figure 3.**
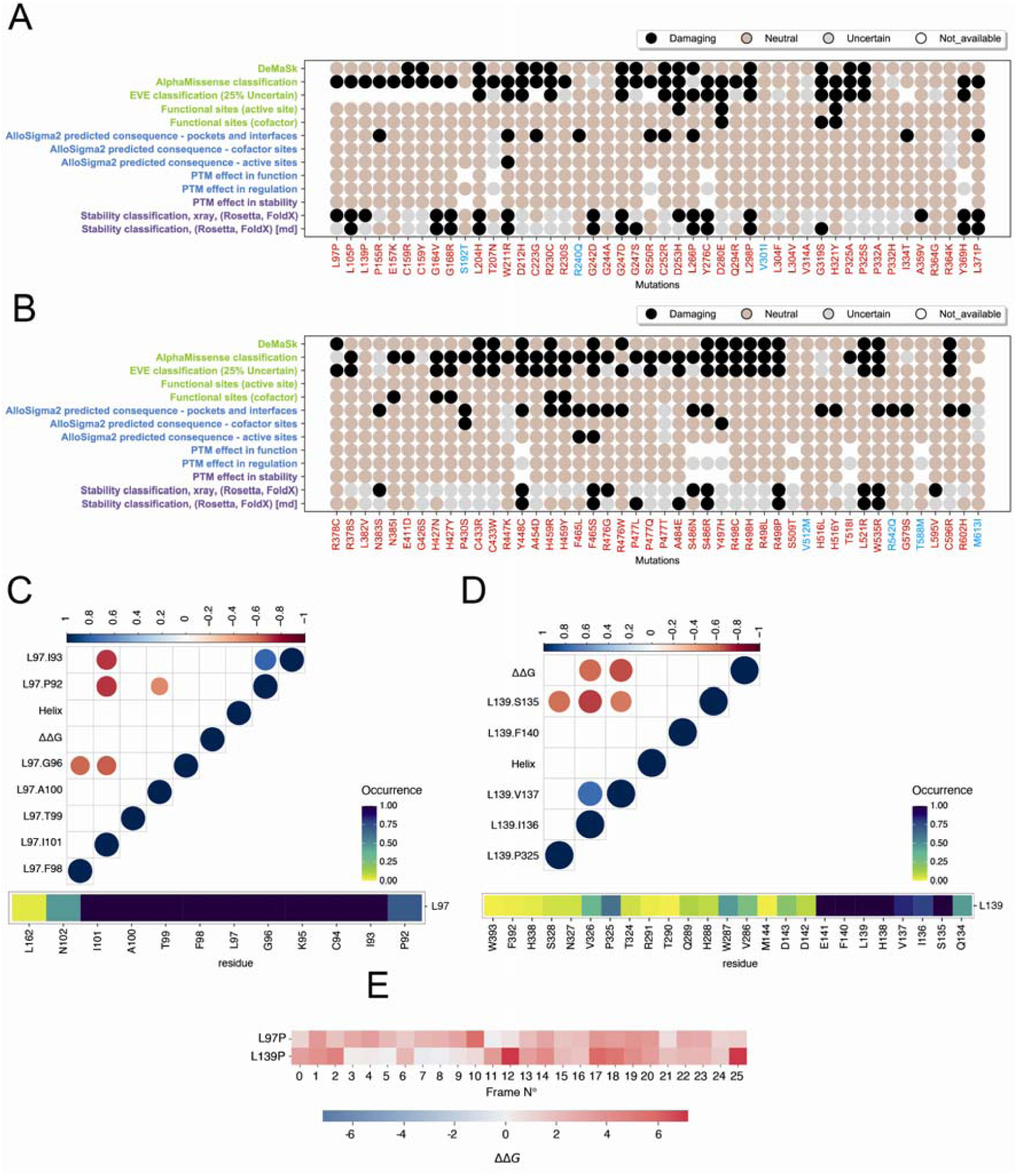
Analysis of likely pathogenic and likely benign variants of ASM reported in ClinVar. A-B)The dot plots from the downstream analysis toolkit of MAVISp illustrate the link between the predicted pathogenic effect and the different mechanisms at play, as covered by the MAVISp framework. We highlighted the known likely benign and likely pathogenic variants from ClinVar in light blue and black, respectively. C-D) Correlation and contact analysis for the mutation sites L97 (C) and L139 (D). E) Folding ΔΔG as estimated for each representative structure of the MD simulation used for the STABILITY module of the ensemble mode of MAVISp for L97P (C) and L139(P). The results show a conformation-dependent effect for the mutations.

ClinVar likely benign variants are predicted with neutral effects in terms of folding free energy and for any of the other MAVISp layers, confirming their benign effect. The only exceptions are R240Q and R542Q, which have a damaging effect on long-range interactions. When investigating these two variants to evaluate if the long-range effect is a genuine signature, we noticed that R542Q is reported to have a stabilizing effect only on L551, which should be excluded from further analysis due to its proximity to the mutation site. R240Q featured communication to multiple sites, of which P529, Q525, and S583 are false positives due to their location in loops around R240, and the remaining sites did not pass the validation step based on PSN-MD.

MAVISp associated a destabilizing effect due to changes in structural stability to 18 of the likely pathogenic ClinVar variants in both the ensemble and the simple mode of MAVISp (L105P, G164V, G168R, L204H, W211R, G242D, G247D, L266P, Y276C, L298P, Y369H, L371P, Y448C, F465S, R476G, S486R, R498P, L521R and W535R). Other four variants (G247S, G319S, P477L, A484E) were classified from uncertain in the simple mode to destabilizing in the ensemble mode, supporting the effect of these variants on structural stability. G247D effects on protein stability are confirmed by the dramatic reduction of the protein half-life for this variant^46^.

On the other hand, when we applied the MAVISp ensemble mode, eight variants were reclassified from destabilizing to uncertain (L97P, L139P, D253H, N383S, R476G, and L595V) or neutral (A359V and S486N). L97P, L139P, and D253H resulted as uncertain variants for stability in the MAVISp ensemble mode due to different predictions by the two methods applied for the consensus approach. N383S, L595V, and R476G, instead, featured average values close to the cutoff used for damaging mutations, which makes it difficult to assess their effect on the structural integrity of the protein in the absence of experimental data to support it.

In the structures from both simple and ensemble modes, L97 and L139 are situated in the α1 and α3 regions of the saposin domain. Throughout the simulation, L97 consistently engaged in hydrophobic interactions with P92, I93, G96, F98, T99, A100, and I101 (**Figure 3C**). Conversely, L139 interacts in more than 60% of the conformations with S135, I136, V137, F140, and P325 and with low occurrence with V286 (i.e., occurrence < 0.25, **Figure 3D**). Substitutions of residues in helical regions to proline are generally expected to disrupt α-helices^47^. However, for L97P and L139P, FoldX, when applied to the MD ensemble, resulted in average ΔΔG values around 2-3 kcal/mol, whereas Rosetta predicted damaging effects. In the MD simulations, the two α-helices populated partially unfolded states. Nevertheless, we did not identify any strong correlation between folding/unfolding events for the α-helices where the two mutation sites are located and the ΔΔG values (**Figure 3C-D**). On the other hand, we observed that the ΔΔG values have large deviations in the 25 structures used to represent the MD ensemble (**Figure 3E**) and that, in the case of L139, seems to be correlated to local contacts with V137 and I136, suggesting that the low ΔΔG values are found in connection with more loose conformations of the α-helix where a substitution to proline would result in a minor effect.

In the ensemble structures, D253 forms stable hydrogen bonds with the backbone and side-chain atoms of H284 (**Table S4**), similar to what is observed in the structure used for the MAVISp simple mode. Concurrently, H284 participates in a π–π stacking interaction with Y490, possibly positioning the histidine for optimal proton donation during catalysis^5^. The substitution of the aspartate with histidine in position 253 will likely strengthen the existing stacking interactions in most models obtained through the FoldX scan on the MD structures (**see Supplementary Movie S2**), suggesting a neutral effect for this variant.

With respect to the structure employed in simple mode, in 20% of the MD frames, the side chain of N383 is involved in a hydrogen bond with the backbone of N385. In comparison, its backbone, similar to the X-ray structure, interacts with the side chain of Q404 in α5 for more than 70% of the MD frames, respectively. This hydrogen bond provides structural support for α5, along with the hydrophobic interaction between F386 P401 and A402 located on the connector of the helix. (**Table S4**). To better evaluate the changes mediated by the side chain, we analyzed the five models of the mutated variant as obtained by each FoldX calculation for each of the 25 frames of the MD ensemble for a total of 125 structures. We observed that the side-chain mediated hydrogen bond formed with N385 by the wild-type residue was lost upon mutation to serine in position 383. Instead, it is replaced by hydrogen bonds of S383 with the main-chain atoms of G366 and G367 (**see Supplementary Movie S3**), suggesting an overall marginal effect of this mutation on structural stability.

R476, located in the β9 strand, establishes hydrogen bonds with high occurrence (> 65%) via its side chain with the E449 side chain. Moreover, E449 participates in a salt bridge with K183 for nearly the entire duration of the simulation (∼ 90%) (see Table S4). R476G causes the loss of this electrostatic network of interaction. However, further studies with MD simulations of the variant would be needed to evaluate if there are compensatory effects from other charged residues in the proximity of E449.

Lastly, L595 in an α-helix in the C-terminal domain is buried (average SASA 3.3% in the MD simulations). We thus estimated the residues in contact with this site in the MD simulation (**Figure S5**). A Val substitution maintains the intermolecular contacts of the wild-type variant and thus is likely to result in a neutral effect on structural stability **(see Supplementary Movie S4)**.

Moreover, for 25 potentially pathogenic variants, the long-range module of MAVISp identifies a potentially damaging effect (**Figure 3A-B**). We scrutinized these variants to evaluate if the signature underlies genuine long-range communication. We identified several mutation sites with predicted allosteric effects that are false positive since the response residue was near the mutation site, or no communication paths were found with the PSN-MD approach. Only W211R, L371P F465L/S, and H459R/Y passed the validation step (**Table S5**). However, we identified only paths at very short distances or circularly connecting back to residues in the proximity of W211.These paths are unlikely to be underlying genuine long-range effect paths to each of its predicted response sites (**Table S5**). W211R also affects the structural stability of the protein, which might dominate with respect to the functional allosteric effect (**Figure 3A**). Supporting its functional importance, W211R causes a dramatic reduction of ASM activity in experimental assays^48^. On the other hand, H459 and L371 were found to be connected to the distal residue P582 (**Table S5**). F465 was connected to E390, near pocket 5 and P332, suggesting a possible long-range effect on the catalytic site (**Table S5**).

C159R/Y, C223G/S/Y, C252R, C433R/W, and C596R are variants reported as Likely Pathogenic in ClinVar, affecting the formation of one of the disulfide bridges of the protein. The results are supported by experimental assays that reported the ASM C159R^49^ and C223S^39^ variants with a residual catalytic activity of 8% and 4%, respectively.

### Variants of ASM reported with conflicting evidence under the lens of the MAVISp framework

48 variants have been reported with conflicting evidence in ClinVar (**Figure 1C**). We thus wonder if the integration of the pathogenicity scores and MAVISp assessment could shed light on their effects.

We identified 11 variants (i.e., N110S, R113C/H, P184L, R202H, R203H, R296Q, V314M, A487V, E517V, V555I) for which most of the pathogenicity scores and all the MAVISp layers used in the study agreed on a neutral effect (**Figure 4A-B**). Of these, experimental data confirmed a mild effect on the catalytic activity (50-70% of residual activity) for A487V and V314M (**Table S3**), strongly supporting that these variants alone could be connected to benign or very mild phenotypes^48,50^. On the contrary, R202H was shown with a drastic reduction of activity (< 5 %, **Table S3**)^39^, and further experimental and structural studies would be needed to understand its effect. At the structural level, it is located at a distal site from the catalytic site and the region of interaction with the membrane. We ruled out allosteric effects at the catalytic site according to MAVISp analysis (**Figure 4A-B**). Nevertheless, it is to be kept in mind that mutations of a residue to histidine could cause structural changes that depend on the pH condition due to the unique pKa range of histidine^51^, and additional simulations of the mutant variant would be needed for a better mechanistic understanding of the effect of this mutation.

**Figure 4.**
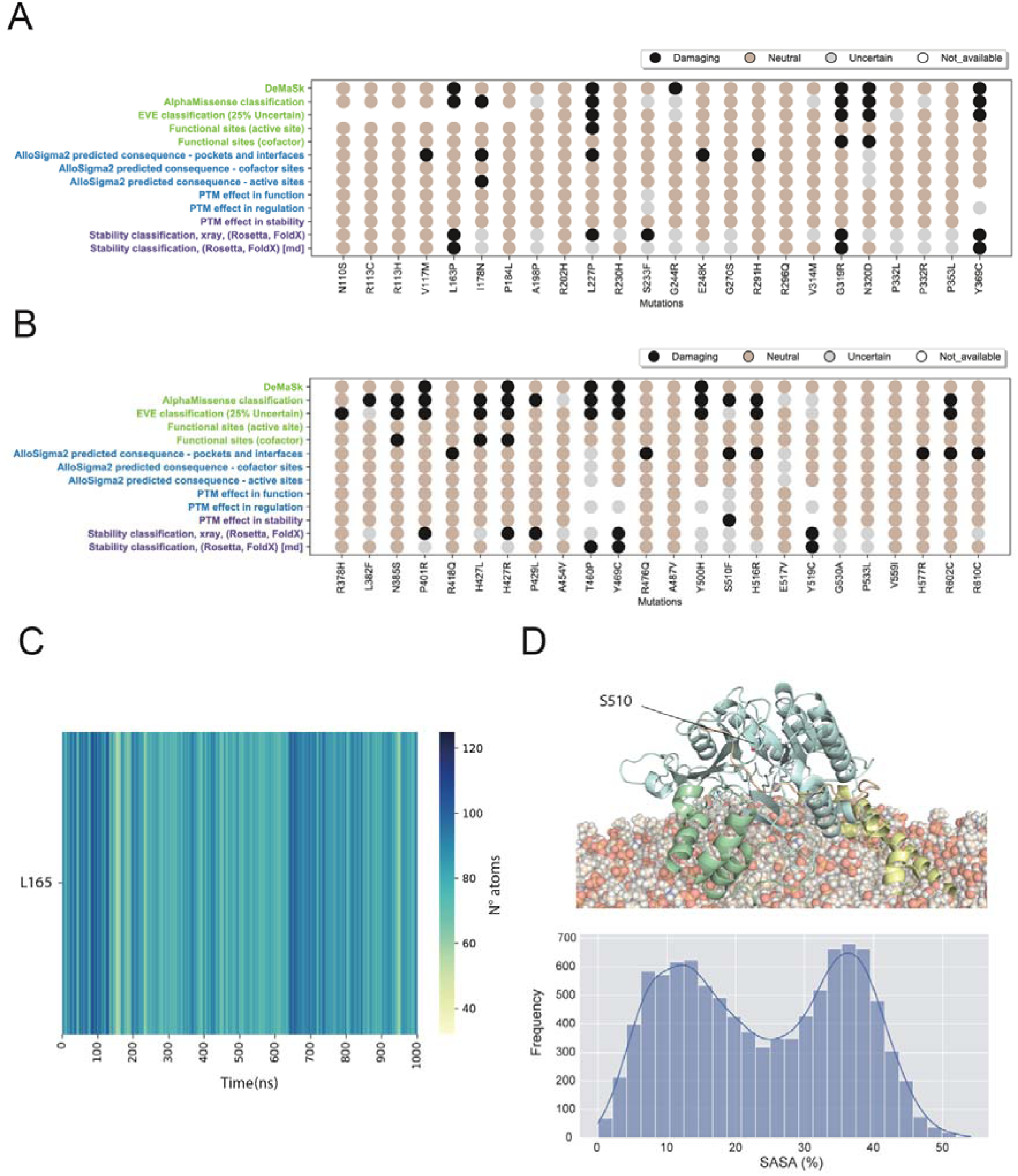
Analysis of ASM variants reported with conflicting evidence in ClinVar. A-B) The dot plots from the downstream analysis toolkit of MAVISp illustrate the link between the predicted pathogenic effect and the different mechanisms at play, as covered by the MAVISp framework. C) The heatmap illustrates the contacts with lipids in the MD simulation for L165. D) S501 is an ASM phosphorylation site that has also been reported mutated in ClinVar located in a region of the protein not involved in membrane association (upper panel. S501 is a buried site in the X-ray structure. Dynamics around the protein native state promote conformational changes that allow the site to be solvent-exposed, as illustrated by the distribution of the relative side chain solvent accessibility of the residue in the MD simulation (lower panel).

On the other hand, we identified a group of potentially pathogenic variants, integrating the results from the pathogenicity scores and the MAVISp modules (**Figure 4A-B**), which we investigated in more detail below.

N320D, N385S, and H427R are associated with alterations at the cofactor binding site, according to the MAVISp assessment. H427 and N320 are Zn^2+^-coordinating residues, whereas N385S is the second sphere of coordination of the Zn^2+^ ion through interactions with N320. The hydrogen bond between N320 and N385 occurs only in less than 2% of the frames of the MD trajectory (**Table S4**). Experimental data identified a detrimental effect on the catalytic activity for N385S (residual activity < 5%, **Table S3**)^52^, supporting the MAVISp prediction. In the case of H427R, a conformation-dependent effect on the folding free energy should also be considered since the predicted effect for the variant changes from highly destabilizing in the MAVISp simple mode to uncertain for stability in the ensemble mode (**Figure 4A-B**).

The MAVISp framework identified L163P, Y369C, T460P, and Y469C as potentially pathogenic variants damaging stability (**Figure 4A-B**). Of note, L163P alters the structural stability and could modify the interaction with the membrane according to the MD simulation. Indeed, throughout the simulation, L163 is in contact with the membrane (**Figure 4C**). This mechanism is relevant because when the ASM interaction with the membrane is altered or lost, the protein is more prone to degradation by cathepsins^53,54^. G319R featured mixed effects with alterations of the folding free energy and an effect on the Zn^2+^-binding (**Figure 4A-B**) as a part of the second coordination sphere through its backbone with H427 (see OSF repository). G319R is known to dramatically decrease the catalytic activity of ASM (**Table S3**)^39^. Our results suggest that the effect is triggered by a higher propensity for misfolding and degradation in this variant more than a direct effect on the catalytic activity (**Figure 4A-B**).

Furthermore, the possible pathogenic effect of I178N, L227P, H516R, and R602C seems to be related to long-range changes according to the analysis in the MAVISp simple mode. We thus investigated them further. Through visual inspection of the 3D structure, we noticed that the response sites W439 for I178 should be considered a false positive. Similarly, for L227P with its response site D214 and R602C with E471. We did not identify communication paths connecting the mutation sites to the remaining response sites with the PSN-MD approach.

S510F pathogenic effect is related to alterations connected to phosphorylation, as evaluated by the PTM module of MAVISp. Interestingly, the residue has a low solvent accessibility in the X-ray structure used in the simple mode, resulting in an uncertain prediction for regulation. In the MAVISp ensemble mode, the solvent accessibility increases (**Figure 4D-E**), suggesting an effect on the regulation of the protein if the phosphorylation is altered. S510 is phosphorylated by PKC-δ kinase^55^. The phosphorylation at S510 is important for ASM activation and translocation to the plasma membrane upon phorbol ester or UV light stimulation ^55^. We also predicted an effect on structural stability due to the removal of the phosphorylation and introduction of a phenylalanine. In particular, the phosphorylation itself is not predicted with a marked effect on the stability of the protein. Still, the replacement with S510 could induce a more destabilizing effect, according to FoldX. Rosetta-based calculations do not provide strong support for this prediction, suggesting that the main effect of the lack of phosphorylation could be in connection to ASM activation and localization.

### Variants of uncertain significance in ASM predicted with damaging effects by MAVISp

Next, we aimed to apply MAVISp to shed light on potentially damaging variants among the ones classified as VUS in ClinVar for a total of 113 scrutinized variants (**Figure 5 A-C**).

**Figure 5.**
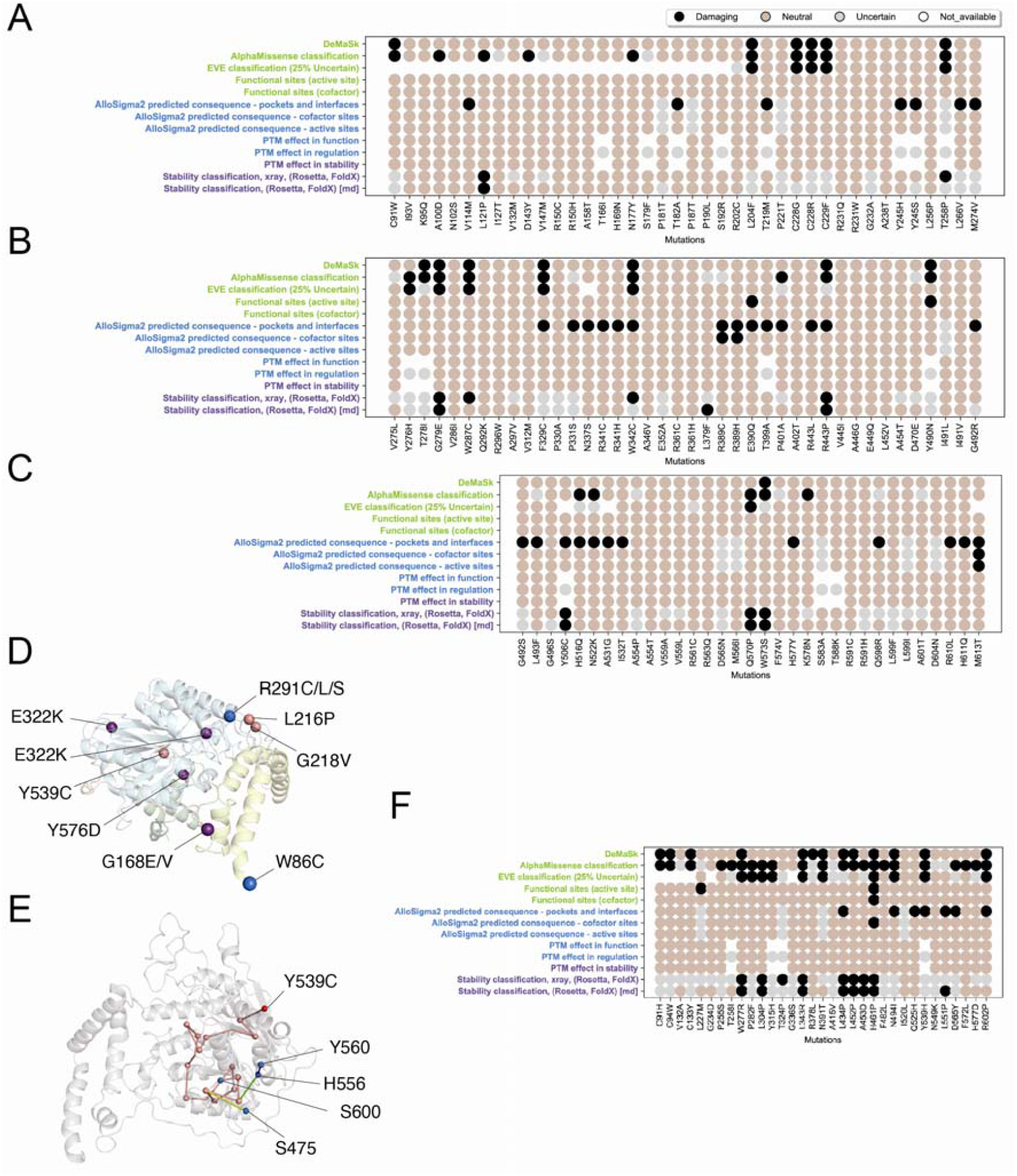
Analysis of ASM VUS as reported by ClinVar, variants identified only in cancer samples and variants curated from the literature. A-C and E) The dot plots from the downstream analysis toolkit of MAVISp illustrate the link between the predicted pathogenic effect and the different mechanisms at play, as covered by the MAVISp framework. The variants reported as VUS in ClinVar (A-C) or curated from literature (F) are illustrated, whereas the variants identified in cancer samples can be found in Figure S6. D) The variant sites found in cancer samples with predicted pathogenic effects and a related MAVISp layer are shown as spheres centered on the Cα atoms of the corresponding residues. Variants are color-coded based on their predicted impact: blue for long-range effects, violet for stability effects, and salmon for variants influencing both long-range and stability. E) The paths of communication to the response sites (in blue) for the mutation site Y539C (in red) are reported with the nodes represented by spheres centered on the Cα atoms of the residues and the edges as cylinders. The paths have been calculated from an atomic contact PSN based on the interaction strength and occurrence in the MD simulation.

At first, we focused on the predicted pathogenic variants by at least one of the three pathogenicity predictors used in the analysis and for which we identified at least one damaging effect by MAVISp. In the case of effects on structural stability, we include the results that were confirmed by the MAVISp ensemble mode.

L121P, G279E, Q570P, and W573S resulted in effects on the structural stability. R443P has predicted effects on both structural stability and long-range. Nevertheless, we noticed that the response sites predicted for an allosteric effect are located near R443, suggesting that the folding free energy altered by this mutation is the most important property.

W342C and H516Q have long-range destabilizing effects on A118 and V114, which we evaluated using the PSN-MD approach and path analysis. P401A has several distal destabilizing effects on different interfaces. After the validation through the PSN-MD step, none of the mentioned variants were retained as no shortest path was found towards the response sites identified by the MAVISp LONG RANGE module.

C91W, C228G/R, and C229F are VUS, which could damage the ASM structure by losing a disulfide bond.

The N522K variant is located at one of the known N-glycosylation sites, abolishing this post-translational modification. The same is observed for the VUS N177Y and N337S. N522K was also predicted with distal effect by the LONG_RANGE module of MAVISp. After a visual inspection of the location of the response sites with respect to N522, we noticed that they can be classified as false positive since it is in proximity to the asparagine residue.

Furthermore, L379F and Y506C are predicted with effects on protein stability by the MAVISp framework. L379F resulted experimentally in approximately 50% residual activity with respect to the wild-type enzyme (**Table S3**)^33^. Looking at the specific values of ΔΔG values, we noticed that they are in the range of 3-3.5 kcal/mol, i.e., close to the cutoff defined based on the relationship between changes in folding free energy and propensity to degradation as estimated by cellular assays^36^.

### Variants only identified in cancer samples

We then focused on the 156 variants reported in COSMIC and cBiPortal as interesting cases to understand the possible damaging effects of ASM at the somatic level in cancer mechanisms (**Figure S6**).

The majority of the cancer variants are predicted with neutral effects by the MAVISp modules, and the pathogenicity scores align with the notion that many of the mutations that accumulate in cancer samples have neutral and passenger effects.

We thus selected the group of mutations predicted pathogenic by at least two of the methods included in the MAVISp framework and for which we could associate a mechanistic indicator (**Figure 5D**) for a total of 12 damaging variants. In detail, G168E/V, E322K, G504E, and Y576D are predicted to be pathogenic due to an alteration of structural stability. W86C and R291C/L/S are predicted with long-term damaging effects using the MAVISp workflow based on Allosigma2. We further investigated removing false positives and in the MD simulations with the PSN-MD approach. As a result, R291C/L/S was predicted to have destabilizing effects and a confirmed communication path to P181 (**Table S5**).

From the analysis of the mutational data of 31 cancer types from The Cancer Genome Atlas^56–58^ (24071849), we found SMPD1 missense mutations in 18 cancer datasets and only one variant R291S in colon adenocarcinoma (TCGA-COAD).

L216P, G218V and Y539C are predicted with both effects on stability and distal effects to ASM solvent-exposed pockets. In the validation phase for LONG-RANGE interactions, L216P and G218V were identified as false positives and subsequently excluded from the PSN analysis. On the other hand, Y539C was predicted with a stabilizing effect on its response sites, as attested by the occurrence of multiple communication paths (**TableS5**). Specifically, Y539C exerted its effect on solvent-accessible pockets for membrane interactions to the response points S475, H556, Y560 (pocket 7), and S600 (pocket 15) (**Figure 5E**). In addition, C94Y and C586R, found in cBioPortal and COSMIC, respectively, caused a loss of one of the disulfide bridges of the protein with a potentially damaging effect.

### Variants from literature curation not reported in cancer databases or ClinVar

33 variants emerged only from literature mining (**Figure 1B**, **Figure 5F**). These variants were not reported or correctly annotated in ClinVar at the time of the analysis (22nd June 2023), and neither are likely to be somatic variants in cancer since we could not find them in CbioPortal or COSMIC. Among those, C91H, C94W, and C133Y resulted in the abolishment of disulfide bridges. The damaging effect is supported by experimental data showing that C133Y abolished the activity of ASM^59^.

Furthermore, we identified W277R, L304P, L343R, L434P, L452P, and A453D as potentially pathogenic through changes in structural stability (**Figure 5F**). L434P seems to have long-range effects from the analysis with the MAVISp simple mode, but the PSN-MD approach does not confirm this. H461P has a strong effect, compromising at the same time the protein stability, the proximity of the active site and the binding to the Zn^2+^ ion. On the contrary, according to the MD simulation, the F572L variant is in a residue placed on an α-helix near the membrane but not in contact with it. The variant F572L at this site was observed to cause a dramatic reduction of the protein half-life, suggesting that the mutated form of ASM could be rapidly degraded and account for the severe clinical phenotype observed^46^. According to the free energy calculations used in this study, the variant is predicted with a neutral effect on folding free energy and tiny changes in kcal/mol using the experimental structure or different conformations extracted from the molecular dynamic simulation.

N494I, Q525H, Y539H, D556Y, and R602P have predicted long-range effects in the simple mode of MAVISp as the main driving feature for their pathogenic effect. After further validating communication routes among these sites and their predicted response site with the PSN-MD approach, only Y539H resulted in distal stabilizing effects, similar to what was shown in the previous section. An effect exerted by this variant is confirmed experimentally by low residual catalytic activity, e.g., 10.2% with respect to the wild-type form^33^.

### Revision of MAVISp parameters toward a more accurate and less computationally intensive variant interpretation for SMPD1

In light of the results above, we noticed many cases of response sites that are not genuinely underlying a distal effect from the mutation site applying the default cutoffs available in the LONG_RANGE module of MAVISp (e.g., distance between C-alpha atoms higher than 10 Å). This requires additional work to identify the damaging variants and visual inspection of the 3D structure. We thus evaluated if increasing the cutoff could result in a clearer picture of the allosteric mutations and the identification of their response sites. We calculated with the LONG_RANGE module, changing the distance cutoffs to 12 and 15 Å (see OSF repository). We noticed that 15 Å were needed to remove the response sites, which are still close to the mutation sites of interest and likely to get closer through conformational changes. This cutoff does not compromise the possibility of identifying response sites that are underlying communication paths as identified by the PSN-MD approach, and it is thus more suited for future investigation on proteins such as SMPD1.

In addition, we used the data collected for SMPD1 to evaluate if, in future studies, we could replace the results from the computationally intensive Rosetta protocol with the ones from a generative model for folding free energy calculations trained on one of the Rosetta energy functions, i.e., RaSP^60^. In the study where the MAVISp framework was presented, it was shown on a dataset of 35 proteins that using a consensus approach for the stability assessment based on RaSP and FoldX resulted in more uncertain variants for stability than the consensus approach based on FoldX and cartesian2020 and the ref2015 energy function^18^. The results here collected on SMPD1 do not confirm the same trend, and we noticed minor differences in the number of variants predicted as uncertain for stability, suggesting that at least for this protein, we could apply RaSP in future studies and variant interpretation (**Figure 6A-F**).

**Figure 6.**
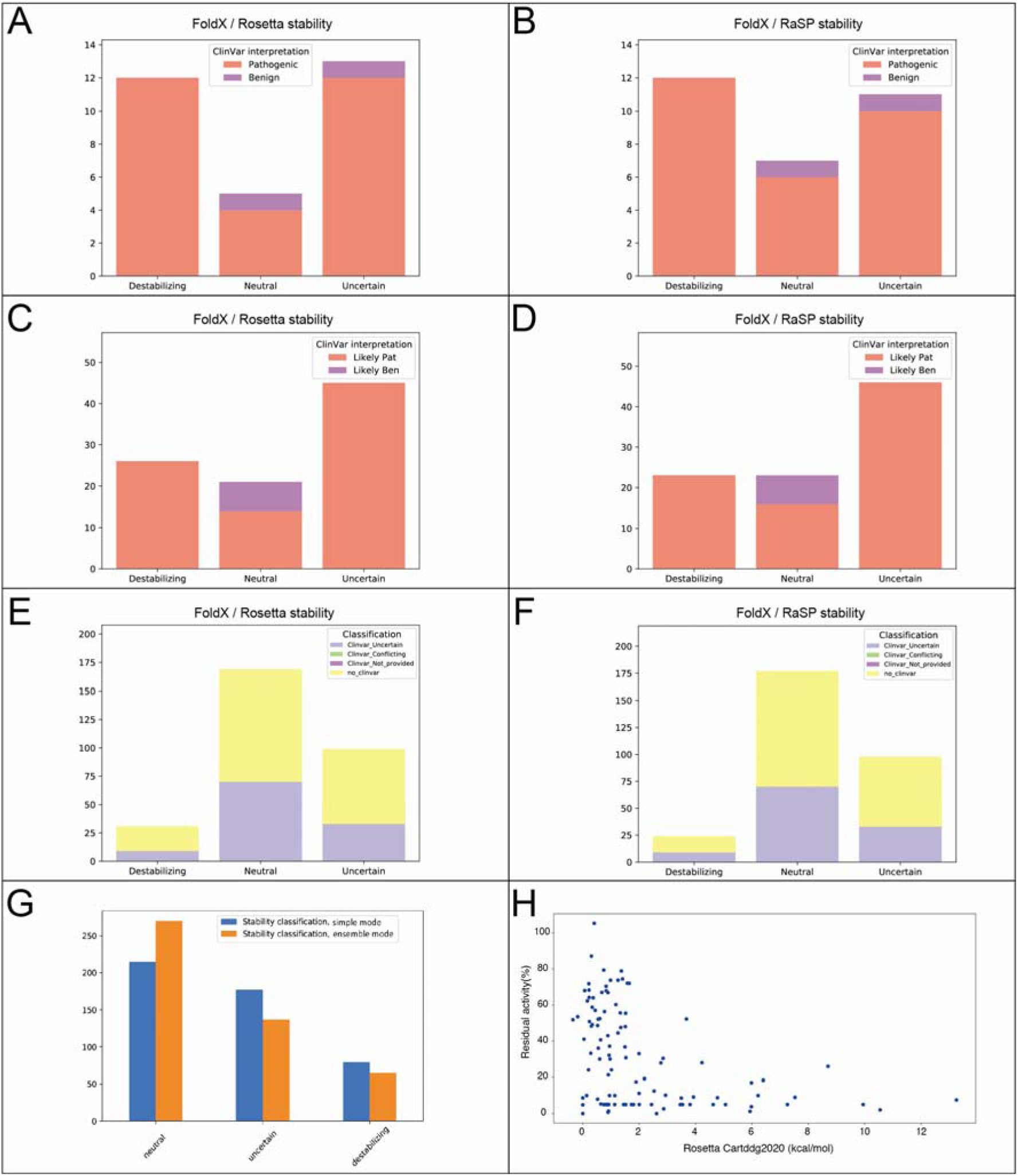
Predicted changes in stability can be used to identify variants with functional effects in ASM. A-G) Comparison of the consensus approach based on FoldX in combination with either Rosetta (A, C, E) or RaSP (B, D, F) for the STABILITY module of MAVISp G) The barplot shows the number of variants for each consequence (i.e., uncertain, neutral or damaging) identified by MAVIsp with the STABILITY module applying the simple or the ensemble mode of the framework. The results show that using the ensemble mode reduces the number of uncertain and damaging variants regarding structural stability. H) The scatterplot shows the distribution of the folding ΔΔG values (predicted by Rosetta) as a function of the residual enzymatic activity of the enzyme. Variants with damaging effects on stability result in residual activity lower than 20%, supporting the predictive power of the approach.

Furthermore, we evaluated if the application of the ensemble mode of MAVISp for prediction of the effects on stability could decrease the number of variants classified as uncertain for this property or if it is rather a way to identify variants that could have damaging effects in a conformation-dependent manner (i.e., as observed above for cases where the classification switches from damaging in the simple mode to uncertain using MD-derived structures) (**Figure 6G**). The results showed a sensible reduction in the number of uncertain variants using the MAVISp ensemble mode with respect to the simple mode. Similarly, we observed a decreased number of predicted destabilizing variants when protein dynamics is taken into account to overcome the limitation of backbone stiffness in FoldX and Rosetta. Overall, our findings support the notion that applying the MAVISp ensemble mode provides a more accurate prediction of the effects of protein variants on structural stability.

### Validation of the selected variants

We curated experimental data from published works on experiments that evaluated the effects of ASM variants on different readouts (i.e., catalytic activity, half-life, and localization). The results are reported in **Table S3.** We then selected from the analyses discussed in the previous sections those variants for which we could assign an effect thanks to the integration of the pathogenicity scores and MAVISp data (**Table 2-3**) and verified the corresponding experimental data.

**Table 2.** ClinVar pathogenic or likely pathogenic variant with an identified mechanistic indicator for their pathogenicity from MAVISp. In the table, we refer to the prediction of pathogenicity with AlphaMissense.

| Variant | ClinVar Classification (review status) | Pathogenicity | Experimental data (residual activity %) | MAVISp interpretation | Structural properties altered |
| --- | --- | --- | --- | --- | --- |
| L105P | 2 | Pathogenic, Likely Pathogenic | < 5 | Damaging | Stability |
| C159R | 2 | Likely Pathogenic | 8.7 | Damaging | Loss of disulfide bridge |
| G168R | 2 | Pathogenic,<br>Likely<br>Pathogenic | 0-20 | Damaging | Stability |
| D280A | 1 | Pathogenic | < 5 | Damaging | Active and<br>Metal-binding<br>sites |
| W211R | 2 | Likely<br>Pathogenic | < 5 | Damaging | Stability |
| C223S | 1 | Pathogenic | 4.8 | Damaging | Loss of disulfide<br>bridge |
| G247D | 1 | Likely<br>Pathogenic | 5-10 | Damaging | Stability |
| D280A | 1 | Pathogenic | < 5 | Damaging | Active and<br>Metal-binding<br>sites |
| A359D | 1 | Pathogenic | 3.8 | Damaging | Stability |
| L363R | 2 | Pathogenic | 1.2 | Damaging | Stability |
| M384I | 1 | Pathogenic | < 5 | Damaging | Stability |
| Y448C | 2 | Pathogenic,<br>Likely<br>Pathogenic | <5, 28 | Damaging | Stability |
| Y469S | 1 | Pathogenic | < 5 | Damaging | Stability |
| P477L | 2 | Pathogenic,<br>Likely<br>Pathogenic | 9.4 | Damaging | Stability |
| S486R | 2 | Pathogenic,<br>Likely<br>Pathogenic | 9.4,9 | Damaging | Stability |
| A484E | 2 | Pathogenic,<br>Likely<br>Pathogenic | < 5 | Damaging | Stability |
| R498P | 1 | Likely<br>Pathogenic | 8.8 | Damaging | Stability |

We noticed that most of the variants predicted with ΔΔG values higher than 3 kcal/mol correspond to residual activities lower than 20 % with respect to the wild-type variant (**Figure 6H**). Prompted by this observation, we partitioned the dataset into two sample sets of residual activity values, using the 3 kcal/mol threshold used to separate destabilizing and other mutaitons (see Methods). We then applied Kolmogorov-Smirnov test between the two sample sets with a resulting p-value (4.78e-05) that support our observation. The data overall point to destabilization of the protein architecture, which in turn can promote misfolded variants more prone to be degraded, as one of the mechanisms to further investigate in disease-associated ASM variants.

Overall, we have identified 18 damaging variants and associated mechanistic indicators integrating pathogenicity scores, MAVISp variant interpretation through different structural properties, and experimental data (**Table 2**). In addition, our results suggest that some known ASM variants found in ClinVar could be reassessed according to what is reported in **Table 3**, including VUS or variants with conflicting evidence.

**Table 3.** VUS or variants suggested for reassessment in light of the data collected in this study.

| **Variant** | **ClinVar Classification (review status)** | **Pathogenicity scores** | **Experimental data (% residual activity)** | **MAVISp interpretation** | **Structural properties altered** |
| --- | --- | --- | --- | --- | --- |
| C133Y | from literature curation | Pathogenic | 0 | Damaging | Loss of disulfide bridge |
| E248Q | Pathogenic (1) | Neutral | 30 | Neutral | none |
| G279E | Uncertain Significance (1) | Pathogenic | N.A. | Damaging | Stability |
| V314M | Conflicting (1) | Neutral | 21, 55.3 | Neutral | none |
| V318M | Pathogenic (0) | Uncertain or Neutral | N.A. | Neutral | none |
| G319R | Conflicting(1) | Pathogenic | 7.57 | Damaging | Stability and Metal-binding site |
| N385S | Conflicting (1) | Pathogenic | < 5 | Damaging | Metal-binding site |
| A487V | Conflicting (1) | Neutral | 73.7 | Neutral | none |
| N522K | Uncertain Significance (1) | Pathogenic | N.A. | Damaging | PTM |
| Y539H | from literature curation | Pathogenic | 10.2 | Damaging | Long Range |
| Q570P | Uncertain Significance (1) | Pathogenic | N.A. | Damaging | Stability |
| W573S | Uncertain Significance (2) | Pathogenic | N.A. | Damaging | Stability |

Finally, we used the MAVISp framework with the modules applied above a saturation, profiling all the possible 10560 variants of ASM in both the simple and ensemble modes. This provides a unique resource for future investigation on new variants or re-evaluation of known ones concerning ASM-related disease (see OSF repository). Of these, 1,549 variants were predicted with a clearly damaging signature in terms of structural stability, which, in turn, would compromise the enzyme activity. More broadly, the combination of MAVISp, EVE, and AlphaMissense allowed the identification of additional pathogenic variants with other structural mechanisms at play, such as a damaging effect from removal of a post-translational modification (S510F) or more than 300 variants with alterations at functional sites.

## Methods

### Dataset of variants used in the study

We collected variants from literature starting from two recent publications ^5,17^ and including additional variants reported in Uniprot (Uniprot entry P17405). We retained only missense mutations in the coding region and removed cases of multiple mutations. This dataset is reported in **Table S1**.

We then applied the MAVISp framework to gather other variants found in ASM, including variants reported in ClinVar^21,22^, COSMIC^24,25^, and cBioPortal^23^. The results can be accessed from the MAVISp database (https://services.healthtech.dtu.dk/services/MAVISp-1.0/).

For ClinVar, we referred to the RefSeq ID NP_000534.4. The protein sequence used in this study is numbered according to the same RefSeq ID, comprising 631 amino acids^1^. This reference introduces a shift of +2 amino acids compared with some published positions. This discrepancy arises from an extra leucine-alanine dipeptide within the polymorphic signal peptide.

### Selection of initial structures for the study

In this study, we collected from both the PDB^61^ and PDBREDO databases ^62^ all the available structures of human acid sphingomyelinase (ASM), specifically PDB entries 5I81, 5I85, 5I8R, 5JG8, 5FIC, 5FI9, and 5FIB^5–7^. The structures were evaluated for missing residues, alternative conformational states for residue side chains, and protonation states using pdb4amber available in AmberTools v2.0^63^, PROPKA 3.1^64^, and H++^65^. Ultimately, the X-ray crystallographic structure of ASM in its open conformation (PDB entry: 5I85) was chosen due to its resolution (2.5 Å) and the conformation of H206, which was appropriately oriented towards the Zn^2+^ ions. Nevertheless, we retained the catalytic water present in the active site of the 5I81 structure and removed the phosphocholine molecule for molecular simulations (see below). To ensure proper orientation of our starting structure within the membrane bilayer, we utilized an oriented structure available on the OPM webserver^66^. For the MD simulation, in contrast to the simple mode, we adopted a protonation state of residues that should be predominant at lysosomal pH, with a pH range of 4.5-5.0^5^. In this setting, we treated lysine and arginine as positively charged, whereas aspartate and glutamate as negatively charged. The histidine residues H138, H169, H213, H284, H288, H321, H338, H432, H511, H534, H556, H577, H580, H611 were positively charged, while H210, H427, and H461 were modeled in the Nε2-H tautomeric state and H423, H459, and H516 in the Nδ1-H tautomeric state (**Table S6**).

We aligned our starting structure with the oriented one to obtain the correct orientation, which we then utilized as the starting structure for the subsequent modelling steps. We used Pymol v2.5 for analysis and visualization of the 3D structure.

### Modeling N-glycosylations of ASM

ASM is N-glycosylated at six sites (N86, N175, N335, N395, N503, N520)^5^. However, no glycoproteomic data is available to identify the specific type of N-glycosylation at those sites to the best of our knowledge. Human ASM was expressed from HEK293S *N*-acetylglucosaminyl transferase I-deficient cells (i.e., cells devoid of complex *N*-glycans) cells, and the X-ray crystallographic structure (PDB ID entry 5I81) shows a truncated oligomannose (potentially Man5) *N*-glycan together with truncated *N*-glycans, with only the first or both GlcNAc of the core resolved^5^. The recombinant human ASM was expressed in CHO cells, and the X-ray crystallographic structure (PDB ID entry 5I8R) is nearly identical to 5I81 but includes fragments of *N*-glycans that could potentially be either complex or oligomannose glycans ^5^. We modeled each *N-*glycan as a Man5 oligomannose using the Glycan Reader tool available in CHARMM-GUI^67,68^.

### MAVISp framework

We used all the applicable modules to this case study from the MAVISp framework to analyze the variants for damaging or neutral effects^18^. We employed both the simple and ensemble modes from the framework. In addition, this study resulted in the design of one new module for local effects on functional sites or cofactors. We also introduced new features within MAVISp for the LONG RANGE module so that the effects on sites such as the active site, substrate binding sites, and cofactor binding sites could be considered. The introduction of these modules was officially included in the MAVISp GitHub repository on 01/11/2023 (https://github.com/ELELAB/MAVISp/pull/215) and 09/11/2023 (https://github.com/ELELAB/MAVISp/pull/217).

In the STABILITY module, we used foldx5 and MutateX, along with Rosetta cartddg2020^69^ protocol and ref2015^70^ energy function with RosettaDDGPrediction^71^ estimate of changes in folding free energy. We used a consensus approach among the two methods to classify the variants, as explained in the original publication^18^. This module was executed both in *simple* and *ensemble* modes.

To estimate changes in long-range structural communication, which could underlie allosteric changes, we applied the LONG RANGE module of MAVISp on an allosteric signaling map generated from AlloSigMA 2^30^. The details about the workflow applied are reported in the MAVISp publication ^18^. For the PTM module, we applied a personalized decision logic to classify each mutation ^18^.

For the INTERACTOME, we applied the workflow of Mentha2PDB from MAVISp^18^ using the dataset of Mentha deposited on 17th April 2023. We also verified with PDBminer^72^ that no structures of ASM in complex with interactors were available in PDB.

We used the results from DeMaSk^20^, EVE^16^, and AlphaMissense^19^ as pathogenicity scores included in the MAVISp framework.

### Molecular dynamics simulations

We performed one-μs all-atom molecular dynamics simulations in explicit solvent for ASM, bound to a mammalian “lysosomal-like” lipid bilayer composition^73^, utilizing the CHARMM36m force field^74^. For the MD simulations, we selected the X-ray crystallographic structure of ASM in its open conformation (PDB entry: 5I85^5^) and modeled five mannose (Man) residues (Man5 *N*-glycans) at each glycosylation site (N88, N177, N337, N397, N505, and N522).

System preparation followed CHARMM-GUI Membrane Builder protocols, involving sequential equilibration steps. At first 10000 steps of energy minimization with the steepest descent method were applied. We then carried out two 250 ps canonical ensemble simulations (NVT) with a one fs integration step, employing the Berendsen thermostat^75^ to reach 310K.

Subsequently, we performed four pressurization steps (250 ps, 500 ps, 500 ps, and five ns). A semi-isotropic pressure control was maintained using the Berendsen barostat with a five ps time constant. The final equilibration phase spanned five ns with a two-fs integration step. Productive simulations employed a two fs time step in an NPT ensemble with the Nose-Hoover thermostat^76^ and Parrinello-Rahman barostat^77^ using one ps and fice ps time constants, respectively. The LINCS algorithm^78^ constrained heavy-atom bonds. We used a cutoff of 12 Å for both Van Der Waals and Coulomb interactions, and the particle-mesh Ewald scheme^79,80^.

### Analysis of the simulations

We used Root Mean Square Fluctuation (RMSF) for each Cα atom of the protein as an index of protein flexibility and used RMSF values averaged over 10-ns time windows. We used NACCESS-based estimates for the relative solvent accessible surface (SASA) for side-chain atoms of the protein residues. We calculated the relative SASA on 10000 frames from the MD simulation. We used PyInteraph2^42^ to calculate electrostatic interactions in the form of hydrogen bonds and salt bridges using default parameters. PyInteraph2 was also employed to obtain an atomic-contact-based Protein Structure Network (PSN) applied to the MD ensemble. We retained the pairs of residues whose sequence distance is higher than 1 for the edge calculations and at a distance lower than 4.5 Å. We used an I_min_ value of 3. Only edges with occurrences higher than 50% of the MD frames were retained in the final PSN, which was weighted according to the interaction strength.

We have used the Dijkstra algorithm^81^ to calculate all simple shortest paths between each pair of nodes of the PSN, as implemented in NetworkX^82^, using the *path_analysis* tool of PyInteraph2. Further, we calculated the normalized node betweenness centrality^83^ as implemented in NetworkX, using the *centrality_analysis* tool of PyInteraph2. All the mutation sites with predicted long-range effects analyzed in this study had normalized betweeness close or equal to 0, and we did not investigate this property further.

We also calculated pair-wise atomic contacts using the *CONtact ANalysis* (*CONAN*) software^84^, applying a *r_cut_* cutoff of 10 Å, *r_inter,_*and *r_high-inter_* values of 4.5 Å.

We calculated the Pearson correlation coefficient and associated p-values for different structural properties and the changes in folding free energy using the *Hmisc*, *PerformanceAnalytics*, and *corrplot* R packages. The results were visualized using a hierarchical clustering approach as a correlogram, applying a p-value cutoff of 0.01.

To investigate the interaction of ASM with the lipid bilayer and quantify the contacts between protein residues and lipid molecules, we computed the number of lipid atoms within a 6 Å radius of the protein residues throughout the MD simulation. We excluded residues from the analysis if all their atoms had lipid contacts in less than 20% of the MD frames.

### Data availability

The aggregated csv data from MAVISp are available at the MAVISp database: https://services.healthtech.dtu.dk/services/MAVISp-1.0/. The MD simulation used for the ensemble mode is available in OSF at https://osf.io/d42zv/. Other accessory data from MAVISp or MD analysis are deposited in the MAVISp extended data OSF repository: https://osf.io/y3p2x/. An overview of the protein and its results from MAVISp is reported in: https://elelab.gitbook.io/mavisp/proteins/smpd1.

## Conclusions

On the one hand, this study allowed us to strengthen, from the methodological point of view, the approach for variant interpretation we recently developed, i.e., MAVISp. Indeed, we introduced new functionalities suited to studying enzymes or metallo-binding proteins. In addition, we identified a better combination of parameters for identifying allosteric mutations and their response sites. The integration of the analysis of communication paths in the MD simulations using a topological network with the results from the allosteric coarse grain model successfully reduced the number of candidates for distal effects. Nevertheless, we cannot rule out that some of the remaining predicted mutations with allosteric effects are genuine and triggered by mechanisms not covered by an atomic contact based PSN. The identification of several mutations in the saposin domain in contact with the membrane also suggests that in the future, a new mechanistic indicator to add to MAVISp could be designed to identify variants that cause structural destabilization and a higher propensity to degradation due to the loss of interactions with the membrane. An approach like this would require that simulations to be also available for the mutant variant but could strengthen the profiling of variant effects in the saposin-domain-containing proteins and not only ASM. Furthermore, we modeled each N-glycan as a Man5 oligomannose in the MD simulation. However, considering the available X-ray crystallographic structures of ASM^5^ and the lack of glycoproteomic analysis, it should be considered that *N*-glycans could be of complex types. A natural future step to elucidate the role of the type of *N*-glycan on the enzyme structure and membrane association would require modeling all *N*-glycans as complex types, such as biantennary digalactosylated systems. Furthermore, it should be noted that tagging with mannose-6-phosphate on the glycans of lysosomal enzymes is responsible for their trafficking to the lysosomes^85^. We omitted mannose-6-phosphate in the design of our glycans based on the previous findings by Zhou et al., which suggested that ASM activity is not influenced by it^5^.

Thanks to the integration of different structure-based methods, machine learning approaches to identify pathogenic mutations and comparison with available experimental data, we here scrutinized and characterized more than 400 variants in the lysosomal Acidic Sphingomyelinase relevant to different diseases, from syndromes related to acid sphingomyelinase deficiency to cancer. Our work also resulted, on one side, in the identification of 18 variants of ASM that have a damaging signature linked to a specific mechanism, including alterations of structural stability (L105P, G168R, W211R, G247D, A359D, L363R, M384I, Y448C, Y469S, P477L, S486R, A484E, R498P, and Y500N) of the active and metal binding sites (D280A), or loss of disulfide bridges (C159R, and C223S).On the other hand, our study, combined with experimental data from the literature, shed light on 11 VUS or variants with conflicting evidence with possible neutral (R296Q, V314M, and A487V) and pathogenic (C133Y, G279E, G319R, N385S, N522K, Y539H, Q570P, and W573S) effects, and pointed out E248Q and V318M for further assessment with respect to the current annotations provided ClinVar.

Changes in structural stabilities are proposed here as a signature of different disease-associated ASM variants, suggesting that one of the next steps is the experimental validation through assays that can dissect these effects. In this context, new advances in the multiplex field^86–88^ could provide a valuable and high-throughput manner to characterize ASM variants for their functional changes. These assays can complement and validate the results from the in-silico saturation scan provided in this work.

Moreover, it should be kept in mind that NPD is a recessive disease, which makes the association between variants and the phenotype challenging to dissect because both alleles have to be considered^13^]. The prediction provided here would imply that the variant is found in both alleles or that the two alleles hold a combination of damaging variants at different sites. In contrast, the presence of only one of the two alleles with a damaging variant could result in milder effects. The knowledge of the individual effects exerted by each variant provided here constitutes the first important step towards understanding their interplay in more complex scenarios.

Altered sphingolipid metabolism has been observed in cancer ^89^ and it has been related to increased tumorigenicity and/or metastatic potential^90–92^. In addition, some cancer therapies have the final effect of inducing ceramide-mediated cell death, but cancer cells evolved escaping mechanisms^93,94^. Recombinant human ASM is a potential treatment for NPD in clinical trials through an approach called enzyme replacement therapy. The latter might also be further explored as an adjunct treatment for cancer therapy^95,96^. This treatment could be especially relevant for individuals carrying variants that compromise ASM structural stability, causing decreased protein levels and where no compensation from other proteins with similar functions is at play. Considering this therapeutical strategy, early and accurate diagnosis is crucial for intervention. The ASM variant atlas provided here can allow us to move a step forward to identify mutations that are potential markers for patients who can benefit from enzyme replacement if combined with other assays and markers.

## Author Contributions (CRediT Classification)

*Conceputalization*: EP *Data Curation*: SS, ML, MT, EP *Formal Analysis*: SS, ML, EP *Funding Acquistion:* EP. *Investigation*: SS, ML, EP *Methodology*: SS, ML, MT, EF, EP *Project administration:* EP *Resources*: EP *Supervision*: EP *Validation*: SS, ML, EP *Visualization*: SS, ML *Writing – Original Draft:* SS, EP. *Writing – Review and Editing*: All the co-authors.

## Acknowledgments

Our research has been supported by Carlsberg Foundation Distinguished Fellowship (CF18-0314), Danmarks Grundforskningsfond (DNRF125), and Novo Nordisk Fonden Bioscience and Basic Biomedicine (NNF20OC0065262) to E.P. group. A EuroHPC Benchmark Access Grant (EHPC-BEN-2023B02-010) supported part of the calculations on Discoverer.

