## Supplementary figures and images for "ASM Variants in the Spotlight: A Structure-Based Atlas for Unraveling Pathogenic Mechanisms in Lysosomal Acid Sphingomyelinase"

### Figure S1

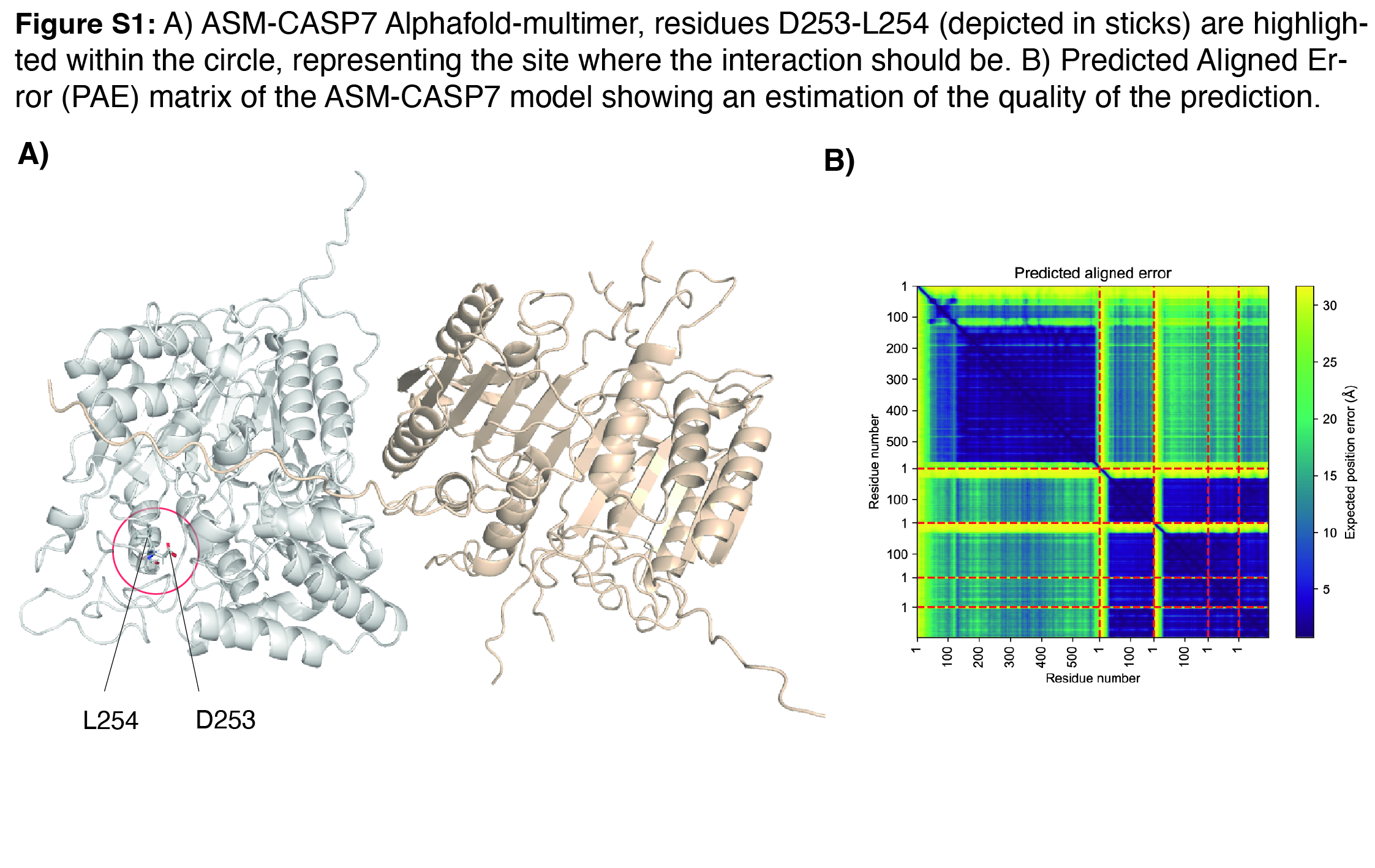

### Figure S2

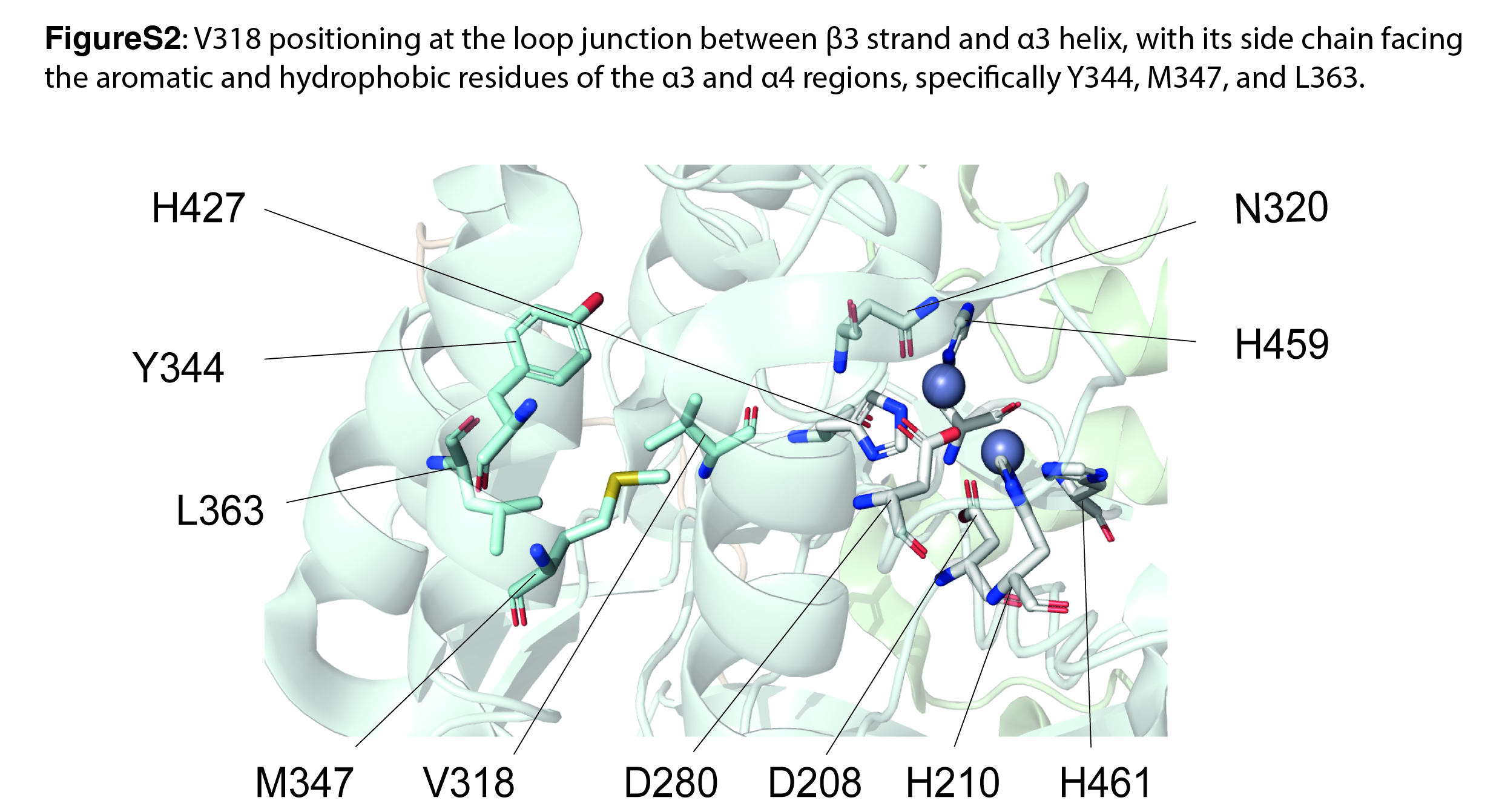

### Figure S3

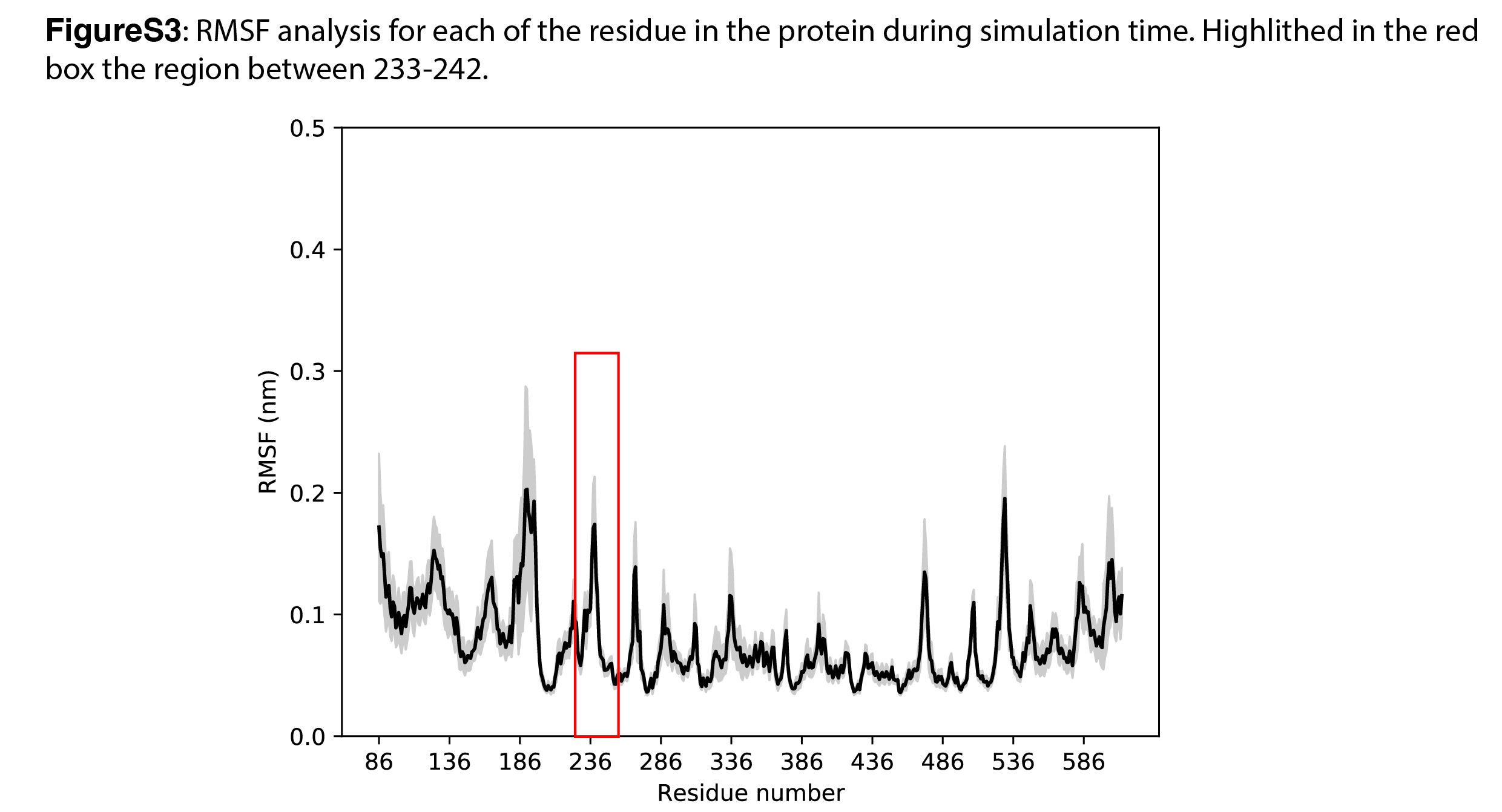

### Figure S4

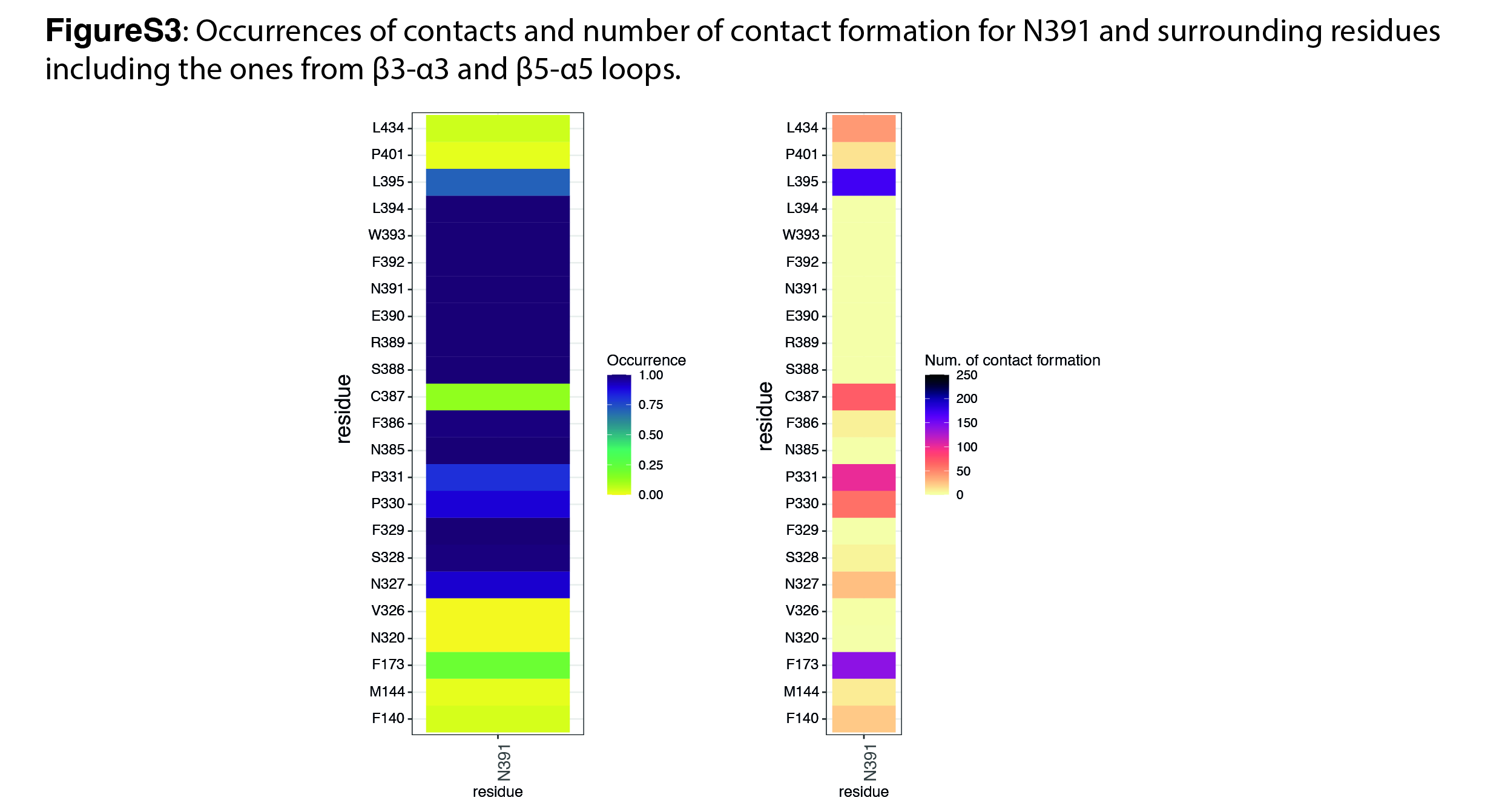

### Figure S5

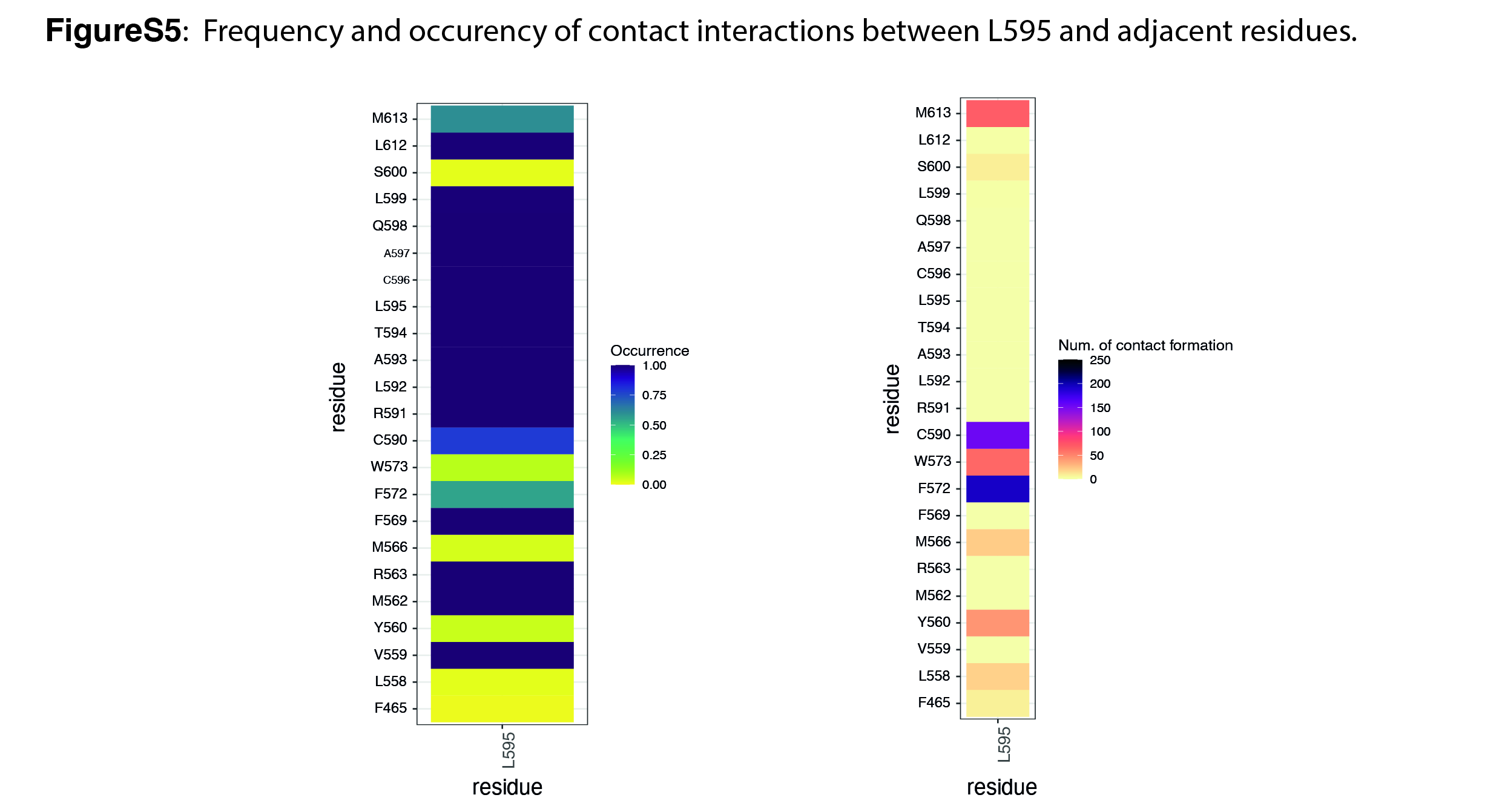

### Figure S6

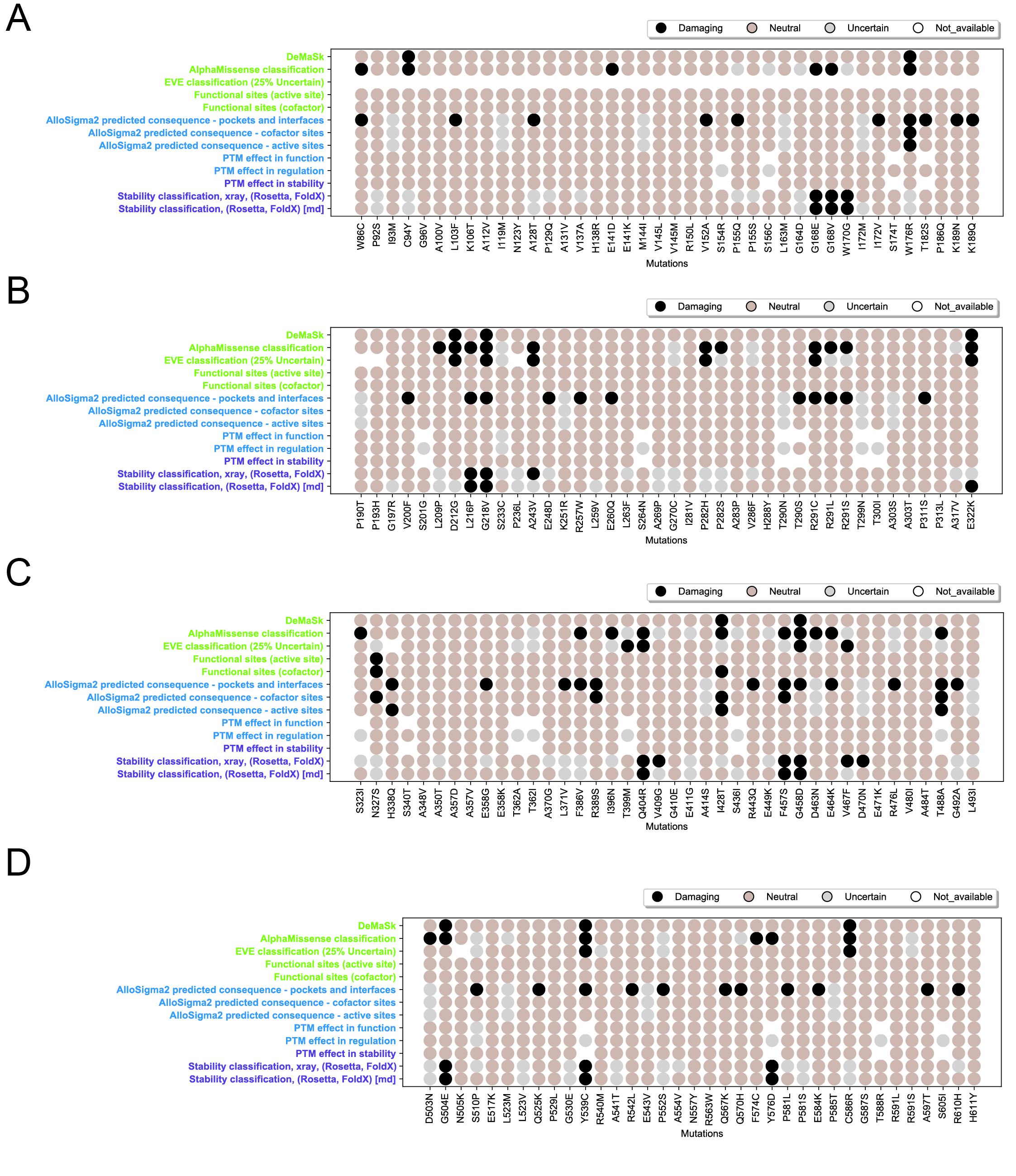

### Supplementary Movie S1

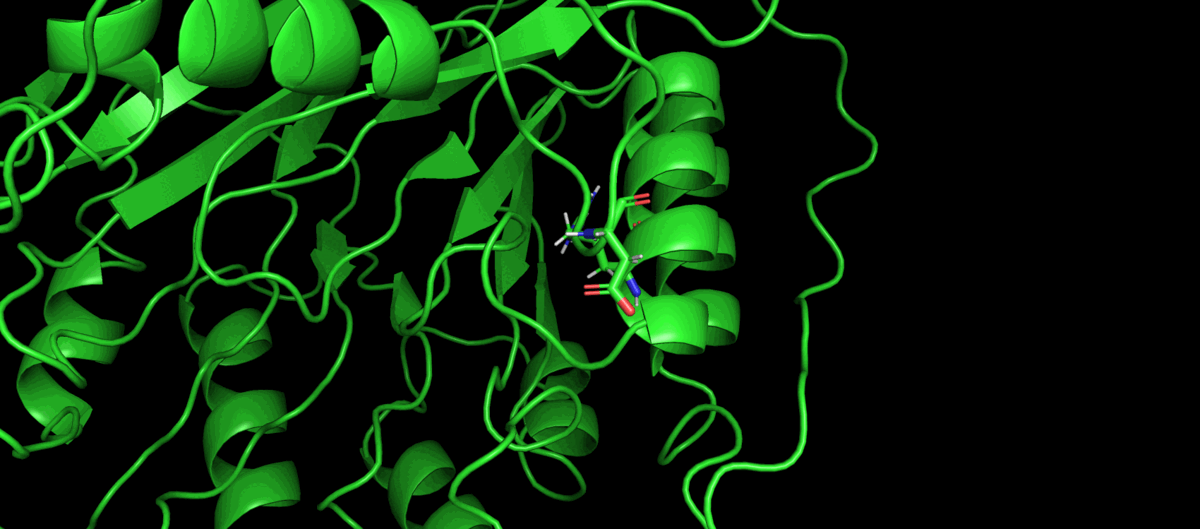

### Supplementary Movie S2

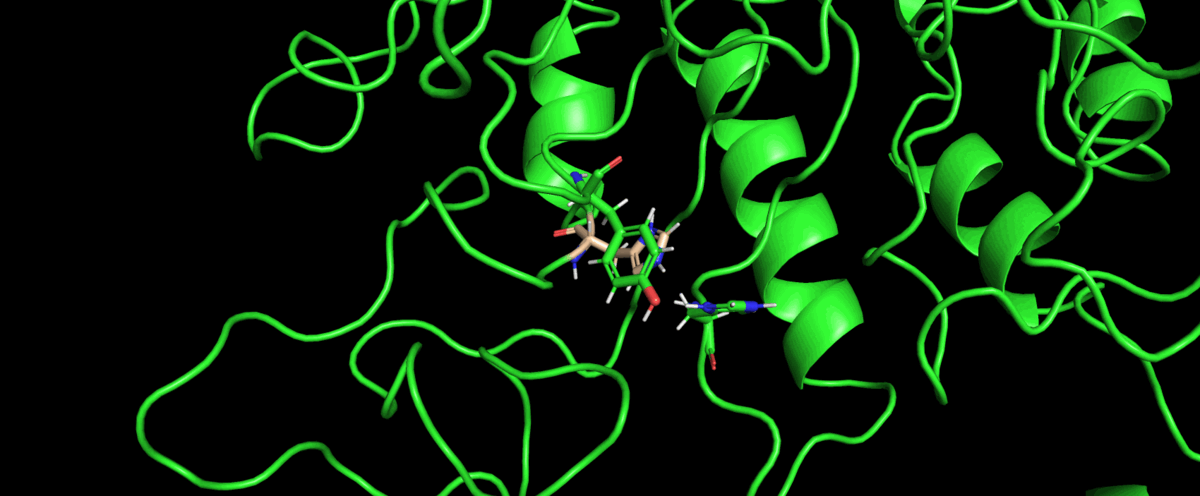

### Supplementary Movie S3

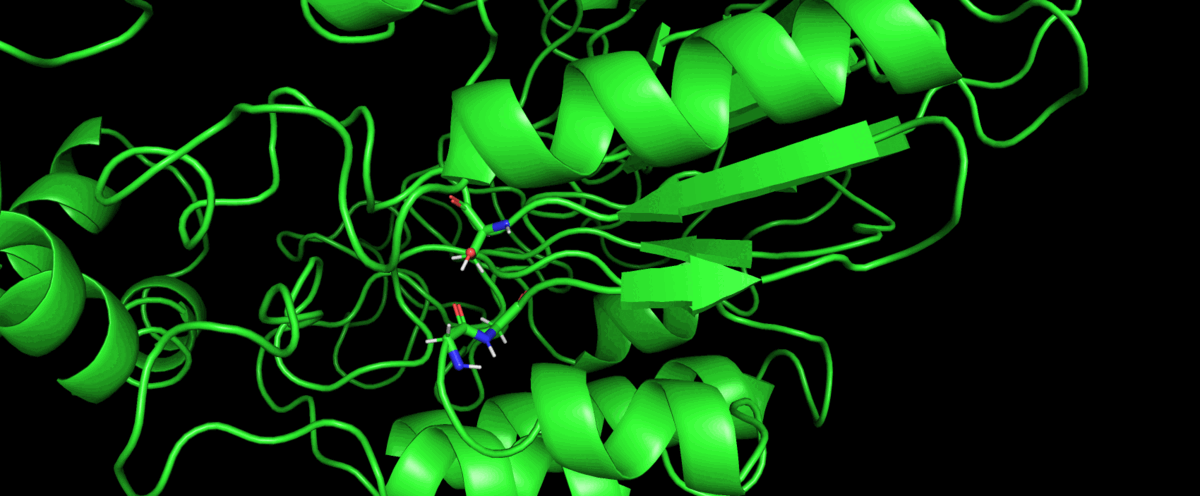

### Supplementary Movie S4

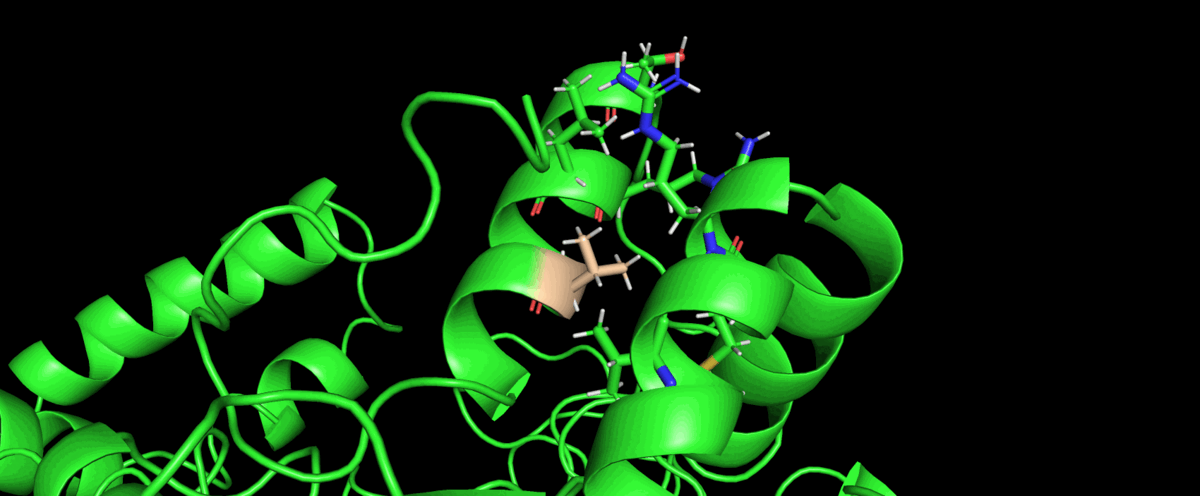
